# Phase-synchronized 40Hz tACS and iTBS effects on gamma oscillations

**DOI:** 10.1101/2025.03.14.643227

**Authors:** Benedikt Glinski, Mohammed Ali Salehinejad, Kuri Takahashi, Asif Jamil, Fatemeh Yavari, Min-Fang Kuo, Michael A. Nitsche

## Abstract

Gamma oscillations play a crucial role in core cognitive functions such as memory processes. Enhancing gamma oscillatory activity, which is reduced in Alzheimer’s Disease, may have therapeutic potential, but effective interventions remain to be determined. Previous studies have shown that phase-synchronized electric and magnetic stimulation boosts brain oscillatory activities at theta, alpha, and delta frequency bands in different ways. The high-frequency gamma frequency band remains to be investigated. This study applies novel noninvasive brain stimulation techniques, namely phase-locked 40-Hz intermittent theta-burst stimulation (iTBS) and transcranial alternating current stimulation (tACS), and explores gamma oscillation changes in the brain. Thirty healthy young participants randomly underwent 40-Hz tACS (1), 40-Hz iTBS (2), two combined interventions (phase-locked iTBS to tACS peak sine wave or tACS trough sine wave) (3-4), and a sham condition (5). The target regions were the left and right dorsolateral prefrontal cortex and were stimulated by simultaneous tACS and iTBS. Gamma oscillatory activities (for 2 hours after intervention) were monitored following each intervention. Our results show that all stimulation protocols enhanced 40-Hz oscillatory power. The iTBS-tACS Peak shows the most significant and stable increase in gamma oscillatory activities (up to 2 hours), followed by 40-Hz tACS and 40-Hz iTBS. 40-Hz tACS and 40-Hz iTBS had the strongest acute effects (up to 30 min) on induced gamma oscillations, while 40-Hz tACS most consistently induced gamma oscillations for up to 2 hours in overall resting EEG data. Phase-synchronizing iTBS with tACS at 40 Hz and the 40 Hz tACS alone targeting the dorsolateral prefrontal cortex, may be a viable approach for inducing and stabilizing gamma oscillatory activity, particularly in conditions where endogenous gamma oscillatory is attenuated, such as Alzheimer’s Disease.

**Registration IDs**:101017716 (CORDIS – European Union-funded project)

## 1 Background

The prevalence of Alzheimer’s disease (AD) is expected to rise threefold by 2050, making AD a serious global health threat [1]. Its growing burden is reflected in rising costs, from 2.8 trillion USD in 2019 to a projected 16.9 trillion USD by 2050 [2]. Besides this current trend, treatment plans for patients mostly focus on symptom management [3]. In addition, new therapeutics targeting amyloid beta (Aβ) plaques, a hallmark of AD, offer limited benefits and risk adverse effects like brain bleeding [4]. This underscores the need for innovative treatments. A recent treatment approach involves non-invasive brain stimulation (NIBS) that includes techniques like transcranial magnetic stimulation (TMS) and transcranial electrical stimulation (tES) that can modulate brain activity.

In TMS, the brain is stimulated by applying single or repetitive magnetic pulses at a specific frequency to the skull. Through electromagnetic induction, magnetic pulses induce an electrical current flow in the underlying brain tissue that leads to changes in electrical and biochemical brain activity [5]. In particular, the repeated application of pulses, repetitive transcranial magnetic stimulation (rTMS), is widely used in neurology, psychiatry, and basic neuroscience to induce long-term potentiation (LTP) and long-term depression (LTD)-like plasticity effects [6] and induce short-lasting changes in oscillatory neural activity in the cerebral cortex [7], [8], [9]. Similar physiological effects are observed for tES interventions, mostly in transcranial direct current stimulation which can induce LTP- and LTD-like plasticity comparable to rTMS. In addition, electrical stimulation with an alternating current, transcranial alternating current stimulation (tACS), interacts with endogenous oscillatory activity by aligning ongoing oscillatory brain activity to the weak exogenous current produced by tACS [10]. This mechanism is called entrainment. The entrainment effects of tACS outlast the acute stimulation phase, leading to longer-lasting changes in oscillatory activity than the short-lasting modulatory effect on oscillations produced by rTMS [11].

Recent evidence suggests that Aβ plaques disrupt plasticity mechanisms and are associated with abnormal oscillatory brain activity in patients compared with healthy controls [12], [13]. Both of these processes can be modulated using NIBS interventions, suggesting their potential as promising strategies for AD treatment. Gamma oscillations play a crucial role in working memory as well as the formation and retention of declarative memory [14], [15], [16]. Previous investigations have shown decreased power of gamma oscillations in the electroencephalogram (EEG) of AD patients [17] and in intracranial recordings of animal models [18], [19]. Reduced gamma oscillations are prominently observed in the hippocampus, visual cortex, auditory, and prefrontal cortex, including the dorsolateral prefrontal cortex (DLPFC) [20], [21], [22]. Specifically, reduced gamma oscillations in the hippocampus are associated with impaired GABAergic interneuron activity, and alleviating these gamma impairments has been shown to restore learning and memory deficits[20]. Additional AD animal studies have reported post-treatment increased Aβ clearance [23], [24]. These findings suggest that mitigating gamma disturbances could represent a novel approach for addressing memory impairments in AD. In this line, modulating 40Hz gamma oscillations enhances Aβ clearance and cognition in preclinical and clinical studies [18], [25], yet challenges remain, including variable patient responses, limited evidence of intervention efficacy and and NIBS protocol optimization [26], [27].

NIBS techniques aiming to interact with 40Hz oscillatory activity could be a promising approach for AD treatment and protocol optimization may improve results. A recent research approach focusing on combining different NIBS techniques showed that the combination of rTMS applied to the peak of the alternating current produced by tACS led to a stronger and longer-lasting increase of mid-range oscillatory power (theta and alpha EEG bands) [28], [29], [30], while for the low-frequency band (i.e., delta) rTMS applied to the trough of the alternating current was most effective [31]. To our knowledge, no study has investigated the effects of this combined stimulation approach for the gamma band. The present study investigated the impact of five NIBS protocols that entrain 40Hz oscillatory activity in the DLPFC of healthy adults. The DLPFC was targeted due to its critical role in working memory and global cognition via gamma oscillations, the disruption of these oscillations in DLPFC by Aβ plaques in Alzheimer’s disease, and its accessibility to NIBS, unlike the hippocampus, making it a common target in NIBS research [32], [33], [34], [35]. We hypothesized that applying intermittent theta burst stimulation (iTBS), a high-frequency rTMS protocol, at the peak of the sinusoidal waveform generated by tACS would produce greater and more sustained increases in 40Hz oscillatory activity compared to iTBS applied at the trough, tACS alone, iTBS alone, or sham stimulation [29], [30]. This hypothesis is supported by findings from non-human primate studies and computational models, which demonstrate that when oscillatory activity becomes synchronized, neurons experience stronger depolarization during the positive phase (peak) of the applied current, whereas a hyperpolarized state occurs during the negative phase (trough) [36], [37], [38].

## 2 Methods

### 2.1. Participants

Thirty healthy subjects (20 females, mean age = 26.54, SD = 3.74) were recruited from the Technical University Dortmund, Ruhr-University Bochum, and the surrounding community. All subjects had normal or corrected to normal eye vision, were right-handed, non-smokers, and had no history of neurological or psychiatric disorders including seizures or epilepsy, CNS medication intake, metal implants, and current pregnancy. To further verify the health status of the subjects, an examination was carried out by a medical doctor before the start of the experiment. Written informed consent was signed by all subjects. The study conformed with the latest version of the Declaration of Helsinki and was approved by the ethics committee of the Leibniz Research Centre for Working Environment and Human Factors (ID: IfADo-196). Participants were monetarily reimbursed and allowed to withdraw from the study at any time.

### 2.2. Intervention

Five interventions with different combinations of iTBS and tACS were tested: (1) *X-tACS Peak* protocol in which the applied iTBS pulses were synchronized to the peak of the target site’s tACS sinusoidal waveform (90° phase from wave onset), (2) *X-tACS Trough* protocol in which these iTBS pulses were synchronized to the trough of the tACS-generated sinusoidal waveform (270° phase from wave onset), (3) the intermittent 40Hz theta burst protocol (iTBS alone) with sham tACS, (4) a 40Hz tACS protocol (tACS alone) with sham iTBS, (5) and a sham protocol (see Figure 1b-e.). We used a modified iTBS protocol (intra-burst interval of 40 Hz instead of 50 Hz), applied at 80% of the individual subjects’ active motor threshold (AMT) four times within a 20-minute time frame starting at 0 minutes, 5 minutes, 10 minutes, and 15 minutes from the beginning of the intervention resulting in 2400 delivered pulses for the whole intervention at each stimulation side. The tACS was delivered utilizing a Starstim 8-channel constant current, battery-powered electric stimulator (Neuroelectrics, Barcelona, Spain). Circular carbon rubber electrodes (2cm radius, 12,57 cm^2^) were used for stimulation. During tACS, current was delivered with a 1mA peak-to-peak amplitude with a 15-second ramp up at the beginning of the stimulation and a 15-second ramp down at the end of the stimulation. Sham tACS was applied by only ramping the current for 15 seconds up and 15 seconds down at the beginning and end of the stimulation respectively. A customized circuit was developed in-house to combine tACS and the modified iTBS protocol by phase-locking iTBS pulses to the tACS waveform. iTBS pulses were delivered using two figure-of-eight-shaped TMS coils. For sham iTBS, two different sham TMS coils (double 70mm pCool-SHAM coil, Mag&More GmbH, Munich, Germany) generated audible clicks at the same intensity as an active coil without inducing an electromagnetic pulse. To minimize the delay between iTBS pulses from the two coils, the TMS devices were connected in series, with TTL signals used for triggering. An oscilloscope confirmed near-zero latency between the pulses. Details of the intervention are in the supplementary material.

**Figure 1:**
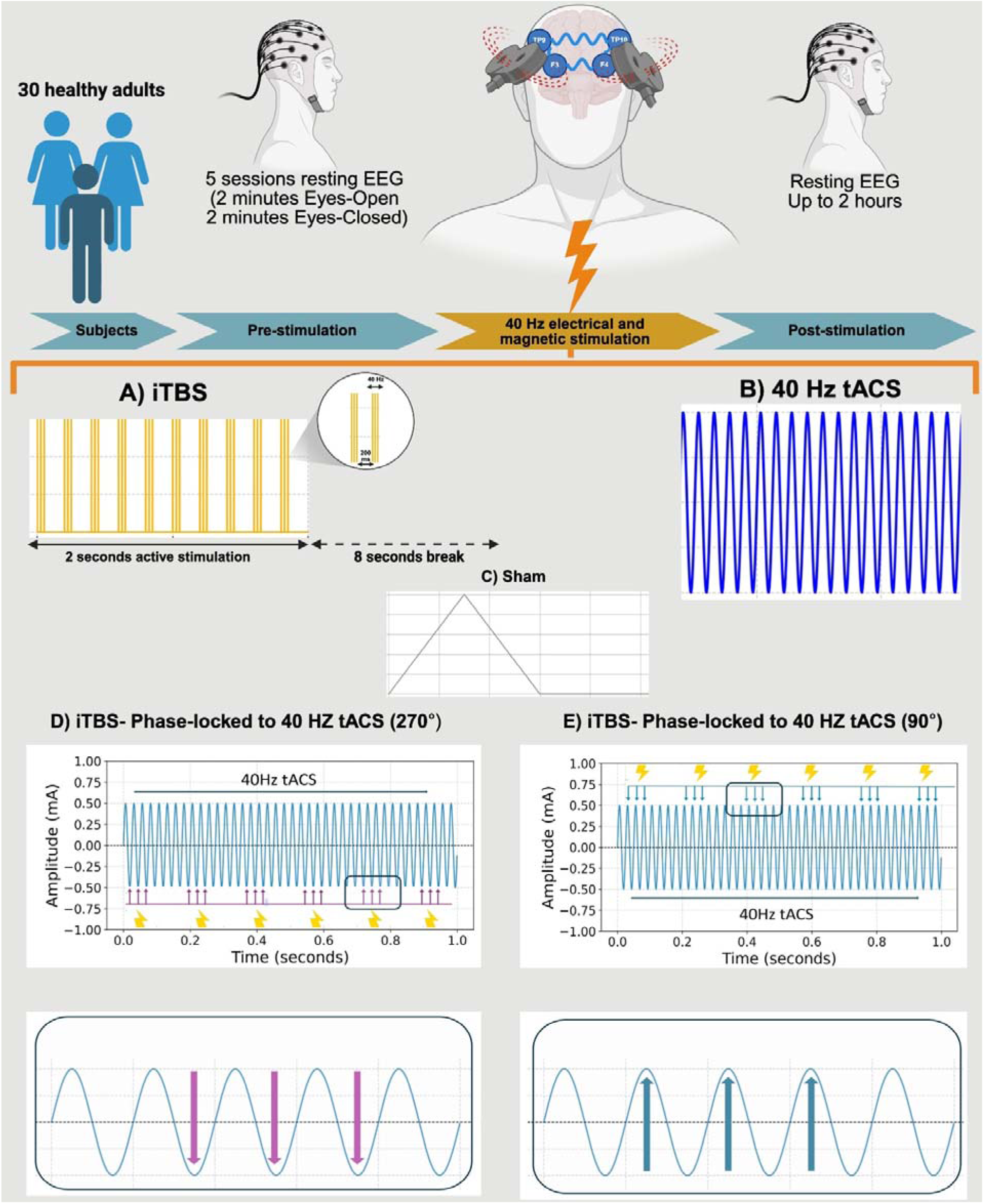
**A)** Graphical representation of the study protocol. **B)** representation of the modified iTBS protocol. The protocol was applied four times in one experimental session. The iTBS cycles started at 0, 5, 10, and 15 minutes leading to a total application of 2400 pulses in one session per stimulation side. Active stimulation in the pattern seen in the representation was applied for 190 seconds per cycle (600 pulses). **C)** Representation of 40Hz tACS with an amplitude of 1 mA peak to peak. The current was delivered for 20 minutes per session. **D)** Representation of the X-tACS Trough protocol. iTBS pulses were phase-locked to the trough (270 degrees) of the sine wave generated by tACS (represented by the arrows). The iTBS intervention was applied as described in b. **D)** Representation of the X-tACS Peak protocol. iTBS pulses were phase-locked to the peak (90 degrees) of the sine wave generated by tACS (represented by the arrows). The iTBS intervention was applied as described in b.

### 2.3. Neuronavigation

To control for movement of the TMS coils during the stimulation, an MR-based 3D-navigation system (LOCALITE GmbH, Germany) was utilized to monitor the orientation and position of the TMS coil to the 3D-head model and fiducials (nasion, left and right ear) of the participants derived from individual anatomical MRI scans. The T1 weighted structural scans were acquired by a Siemens 3T Prisma scanner based on the 3D-MP-RAGE sequence (TR=2600 ms, TE=2.92 ms, flip angle=12°, FoV read=256 mm, FoVphase=87.5%, 176 slices with 1×1×1 mm^3^ resolution).

### 2.4. Resting Electroencephalography

EEG was recorded using a 64-channel EEG System (Bittium, NeurOne, Bittium Corporation, Finland) in a shielded room with stable light conditions. The Ag/AgCl EEG electrodes were attached to the head using high-viscosity electrolyte gel (SuperVisc, Easycap, Herrsching, Germany). All recordings were sampled with a sampling rate of 2000 Hz. The online reference electrode was placed at FCz and the ground electrode was placed at CPz. The EEG impedance was kept below 5kOhms throughout the whole recording. Digital TTL triggers were transmitted to the EEG system by a server PC to indicate the start of a condition (eyes open or eyes closed) during the resting state recordings. For EEG preprocessing, BrainVision Analyzer version 2.2.2 (Brain Products GmbH, Gilching, Germany) was used. The pipeline was adapted from Salehinejad et al. (2021). EEG recordings were bandpass-filtered between 0.1 and 90Hz (zero phase shift Butterworth filters) with a notch filter at 50Hz to remove line noise. All electrodes were referenced to a common average reference montage. The vEOG signal from channel Fp2 was used for the correction of ocular artifacts using the Gratton and Coles method [40] in line with our previous works [29–30]. Channels with artefactual signals were identified and spline-interpolated. Resting state EEG recordings were then segmented into two-second epochs, artifactual epochs containing EEG activity greater than a 100mV peak-to-peak amplitude automatically marked, visually inspected, and removed manually. Artifact-free segments were then transformed from the time to the frequency domain using Fast Fourier transform (Hanning window length 10%, zero padded) and averaged.

### 2.5. Experimental design and procedure

Participants attended 5 experimental sessions (see Figure 1A), all scheduled at the same time of the day (9 am or 1 pm) to control for arousal levels and sleep pressure [39], [41]. Sessions were spaced at least seven days apart to prevent carry-over effects. EEG recordings were conducted in a shielded, soundproof chamber with consistent lighting. Upon arrival, participants were seated in a reclined chair, and a 64-channel EEG cap was fitted according to the 10-10 international electrode placement convention. tACS electrodes were pre-placed over the target regions (F3, F4, TP9, and TP10). After the EEG setup, a vacuum pillow was used to stabilize the head position. The TMS intensity for iTBS was determined in each session separately to account for changes of cortical excitability. Baseline EEG recordings consisted of two minutes each of eyes open (EO) and eyes closed (EC) conditions. During the EO recording, participants fixated on a black cross against a white background at one meter eye distance.

### 2.6. Statistical analysis

Data were analyzed using R version 4.4.0 (R Core Team, 2024) and GraphPad Prism version 9.5.1 (GraphPad Software, Boston, Massachusetts, USA). Physiological data (EEG power spectrum and functional connectivity) were preprocessed and analyzed with Brainvision Analyzer, Fieldtrip, and Brainstorm toolbox [42], [43]. Figures were created using Prism 9.5.1, Microsoft PowerPoint, and MATLAB 2023b utilizing the Fieldtrip toolbox. Details of data analysis (physiology, behavior, blinding, and side effects) are presented in the supplementary material.

## 3 Results

### 3.1. Data overview, blinding, and side effects

No significant differences were observed in participant ratings whether real (X-tACS Peak, X-tACS Trough, tACS, iTBS) or sham stimulation was applied, confirming the effectiveness of blinding (Table S1). The side effects reported immediately and 24 hours post-stimulation are summarized in Table S2. No significant differences were observed in tingling, burning, pain, or skin redness across stimulation conditions. However, a significant effect of stimulation on visual phenomena was detected (Table S3). Post hoc analyses revealed significantly larger visual phenomena were observed for tACS (t = 3.67, p = 0.001), X-tACS Trough (t = 3.54, p = 0.001), and X-tACS Peak (t = 3.61, p = 0.001), while iTBS again showed no significant difference (p = 0.801). Follow-up correlational analyses indicated a significant positive correlation (p = 0.015) with moderate strength (r = 0.45; see Figure S2) between visual phenomena and post-intervention 40Hz oscillatory power in the frontal ROI only in the eyes open condition (EEG sensors: F3, F4, Fz, F1, F2, AF3, AF4, FC3, FC4, F5, F6) specifically for the tACS protocol in the EO condition. This correlation, however, was not significant at the whole-brain level (r = 0.35; p = 0.065). The remaining non-significant correlations between visual phenomena and induced 40 Hz gamma oscillations were as follows: r = 0.09 and r = 0.20 for the X-tACS Peak at the frontal ROI and whole-brain level, respectively, and r = −0.92 and r = 0.07 for the X-tACS Trough.

### 3.2. Stimulation effects on 40Hz oscillatory power

The linear mixed model (LMM) analysis revealed no significant baseline differences between the absolute 40Hz power of the different stimulation protocols in the EO, EC and and overall conditions in the frontal ROI and at the whole brain level (Figure S3). Temporal dynamics of the interventions focusing on the frontal region of interest were analyzed using separate LMMs for the EO and EC and overall recording conditions. The analyses revealed significant main effects of stimulation (EO: F_4,6979_= 43.90, p<0.001; EC: F_4,7111_=17.25, p<0.001; overall: F_4, 7076.7_=16.90, p<0.001), time (EO: F_4,6978_= 85.41, p<0.001; EC: F_4,7111_=121.82, p<0.001; overall: F_4, 7076.2_=97.49, p<0.001), and electrode (EO: F_10,6978_=5.36, p<0.001; EC: F_10,7111_=4.01, p<0.001; overall: F_10, 7076.1_= 8.01, p<0.001), and a significant stimulation × time interaction (EO: F_16,6978_= 5.86, p<0.001; EC F_16,7111_=2.03, p<0.001; overall: F_16, 7076.1_=2.54, p<0.001) (Table 1). Subsequent pairwise comparisons revealed temporal and stimulation-specific effects.

**Table 1:** LMM results: Significant main and interaction effects of stimulation, time and electrode on 40Hz oscillatory power in the frontal ROI.

| Condition | Factor | d.f. | F value | p value | $\eta p^2$ |
| --- | --- | --- | --- | --- | --- |
| **Eyes-open** | stimulation | 4, 6979 | 43.90 | **<0.001** | 0.02 |
|  | time | 4, 6978 | 85.41 | **<0.001** | 0.06 |
|  | electrode | 10, 6978 | 5.36 | **<0.001** | 0.00 |
| | stimulation × time | 16, 6978 | 5.86 | **<0.001** | 0.01 |
| | stimulation × electrode | 40, 6978 | 1.02 | 0.43 | 0.00 |
| | electrode × time | 40, 6978 | 0.65 | 0.96 | 0.00 |
| | stimulation × time × electrode | 160, 6978 | 0.38 | 1.00 | 0.00 |
| **Eyes-closed** | stimulation | 4, 7111 | 17.25 | **<0.001** | 0.00 |
|  | time | 4, 7111 | 121.82 | **<0.001** | 0.06 |
|  | electrode | 10, 7111 | 4.01 | **<0.001** | 0.00 |
| | stimulation × time | 16, 7111 | 2.04 | **<0.01** | 0.00 |
| | stimulation × electrode | 40, 7111 | 2.81 | **<0.001** | 0.02 |
| | electrode × time | 40, 7111 | 0.83 | 0.77 | 0.00 |
| | stimulation × time × electrode | 160, 7111 | 0.41 | 1.00 | 0.00 |
| **Overall (EO+EC)** | stimulation | 4, 7076 | 16.89 | **<0.001** | 0.00 |
|  | time | 4, 7076 | 97.49 | **<0.001** | 0.05 |
|  | electrode | 10, 7076 | 8.01 | **<0.001** | 0.01 |
| | stimulation × time | 16, 7076 | 2.54 | **<0.001** | 0.00 |
| | stimulation × electrode | 40, 7076 | 1.97 | **<0.001** | 0.01 |
| | electrode × time | 40, 7076 | 0.86 | 0.72 | 0.00 |
| | stimulation × time × electrode | 160, 7076 | 0.39 | 1.00 | 0.00 |

#### 3.2.1. 40Hz oscillatory power at frontal ROI

##### 3.2.1.1. Eyes open condition

Compared to baseline, X-tACS Peak, tACS, and iTBS significantly increased 40Hz power during the first 30 minutes post-intervention, (all p<0.001), with no effect for X-tACS Trough, and sham conditions. At the 30-60 minutes time window, all active protocols increased 40Hz power (all p<0.001) and remained significant until 120 minutes post-intervention (all p< 0.001). Sham stimulation did not produce significant changes besides the 90-120 minutes time window when an increased 40Hz power was observed (t=3.08; p=0.010; Table S8). When compared to the sham condition, X-tACS Peak, X-tACS Trough, tACS, and iTBS protocols significantly increased 40Hz oscillatory power during the first 30-minutes post-intervention (all p<0.05). The effects of X-tACS Peak, tACS and iTBS stimulation were not statistically different from each other. Furthermore, X-tACS Peak, tACS and iTBS stimulation protocols had a stronger increase of 40Hz gamma oscillations compared to the X-tACS Trough protocol (all p<0.05).

In the 30-60 minutes time window, all protocols except tACS resulted in a significant increase in 40Hz oscillatory power compared to the sham condition (all p< 0.01). Here, the X-tACS Peak protocol produced the most substantial enhancement, significantly surpassing the effects produced by X-tACS Trough, tACS, and iTBS (all p<0.05). No other significant effects were found. In the 60-90 minutes time window, all protocols significantly increased 40Hz oscillatory power compared to sham (all p<0.001). X-tACS Peak and tACS showed the largest effects, both outperforming iTBS and X-tACS Trough (p<0.001). Finally, during the 90-120 minutes time window, all protocols maintained significantly higher 40Hz oscillatory power compared to sham (all p<0.001). The X-tACS Peak stimulation condition induced a significantly larger increase than both X-tACS Trough and iTBS (all p< 0.05), but not compared to tACS (t = −1.99, p=0.07). Differences between iTBS, X-tACS Trough, and tACS conditions were not statistically significant (Figure 2A). For detailed statistics see Table S7/8 for effects in other frequency bands see Figure S4).

**Figure 2:**
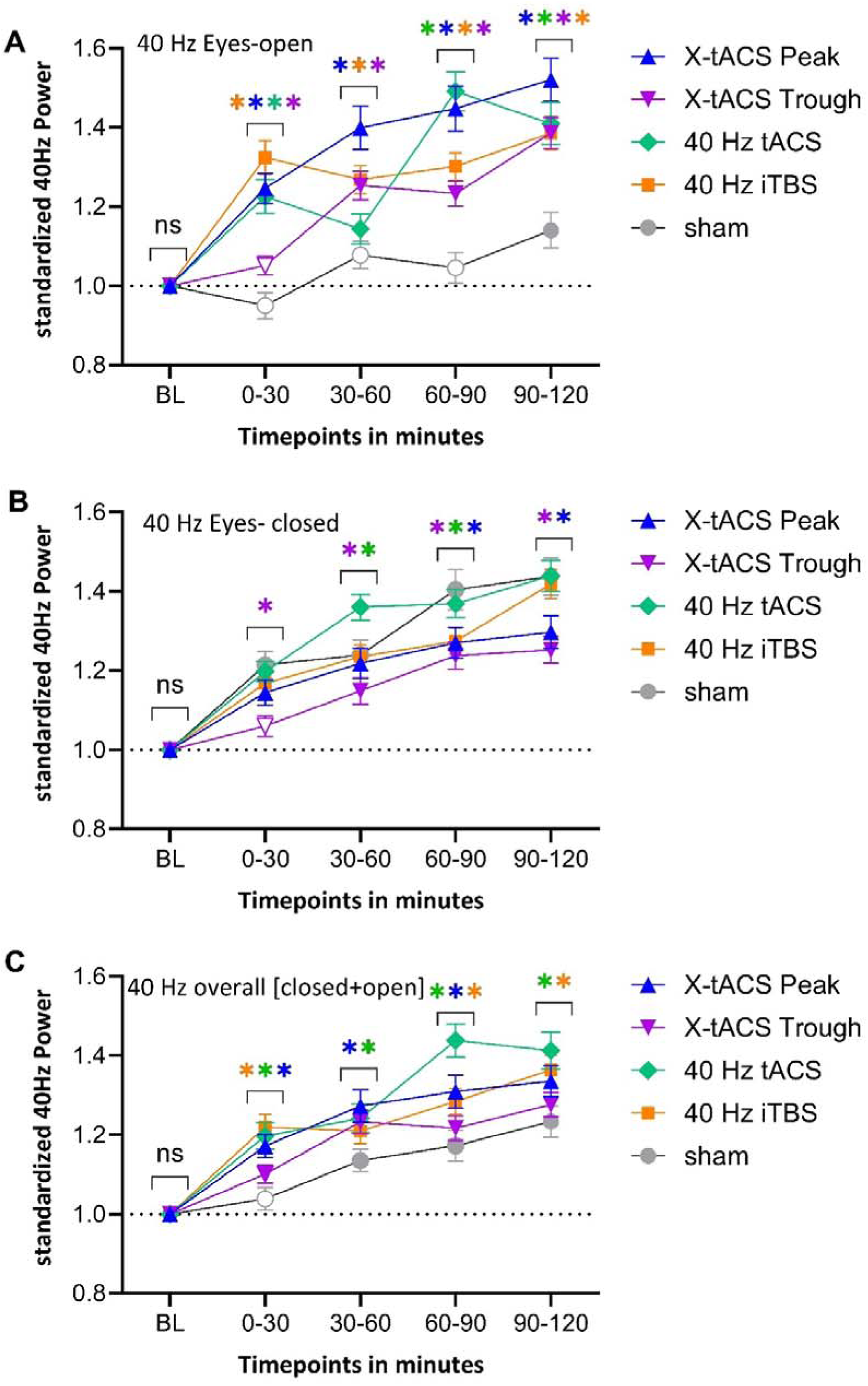
**A,** Relative time-dependent changes in 40Hz power for all stimulation protocols in the frontal ROI, compared to the baseline during **eyes-open** recordings, are shown. **B,** Relative time-dependent changes of 40Hz power in the frontal ROI are shown for all stimulation protocols in relation to the baseline for the **eyes closed** recordings. **C,** Relative time-dependent changes in 40Hz power for all stimulation protocols in the frontal ROI, compared to the baseline during **eyes-open+eyes closed (overall)** recordings, are shown. In a, b and c, filled symbols indicate a significant difference compared to the baseline. Asterisks above the graph indicate a significant difference compared to sham (p<0.05, FDR-corrected). Error bars represent the standard error of the mean.

##### 3.2.1.2. Eyes closed condition

All interventions, including sham, except for X-tACS Trough during the first 30 minutes post-stimulation (t=1.41; p=0.18), significantly increased 40Hz oscillatory power relative to baseline (all p< 0.01). When compared to the sham condition, X-tACS Trough resulted in decreased 40Hz power during the first 30 minutes (t=-3.76, p<0.01). No other significant differences were found compared to the sham condition. Compared to all other active protocols, X-tACS Trough significantly decreased 40Hz oscillatory power (all p< 0.05). In the 30-60 minute time, tACS increased 40Hz power (t=2.57, p=0.03) whereas X-tACS Trough decreased 40Hz power (t=-2.41, p=0.03) when compared to sham. The increasing effect of tACS on 40Hz power in the frontal region was higher than for all other tested protocols (all p<0.05). No other significant effects were observed during this time window.

At the 60-90 minutes post-intervention, all protocols except tACS reduced 40Hz power compared to sham (all p<0.001). The effect of tACS did not differ significantly from sham (t=-1.21; p=0.31) but was significantly different from the X-tACS Trough protocol (t=3.08; p<0.01), which caused the greatest reduction. No other significant differences were observed between the protocols. Finally, during the period of 90-120 minutes, only the combined protocols significantly reduced 40Hz power compared to sham (p <0.05). This decreasing effect of X-tACS Peak and X-tACS Trough was also significant when compared to iTBS and tACS (p<0.05). No significant difference was observed between X-tACS Peak and X-tACS Trough conditions (Figure 2B; Table S9-10; for effects in other frequency bands see Figure S4).

##### 3.2.1.3. Eyes closed + eyes open condition

Compared to baseline, all active stimulation protocols significantly increased 40 Hz gamma oscillation at all time points. The sham condition showed similar increases except during the first 30 minutes. When compared to the sham, in the first 30 minutes post-stimulation, 40 iTBS and tACS, followed by X-tACS Peak, significantly increased 40 Hz gamma oscillations compared to the sham. At 60-90 minutes post-intervention, these same protocols significantly enhanced 40 Hz gamma oscillations. Only tACS and X-tACS Peak enhanced 40 Hz gamma oscillations at 30-60 minutes post-intervention. At 90-120 minutes post-intervention, only 40 iTBS and tACS increased 40 Hz gamma oscillations. 40 Hz tACS was the only stimulation protocol that significantly increased gamma oscillation in all timepoints.

### 3.3. 40Hz oscillatory power at whole-brain level

The effects of stimulation on the whole brain level were investigated using separate LMMs for EO and EC recording conditions. Significant main effects were found for stimulation (EO: F_4,38638_=236.19, p<0.001; EC: F_4,38309_ = 149.64, p<0.001), time (EO: F_4,38637_=470.71, p<0.001; EC: F_4,38309_=737.70, p<0.001), electrode (EO: F_60,38637_=8.94, p<0.001; EC: F_60,38309_=7.01, p<0.001), and significant interactions for stimulation × time (EO: F_4,38637_=29.10, p<0.001; EC: F_4,38309_=17.01, p<0.001) and electrode × stimulation (EO: F_240,38637_=2.19, p<0.001; EC: F_240,38309_=2.22, p<0.001). No other significant effects were found (Table 2).

**Table 2:** LMM results: Significant main and interaction effects of stimulation, time and electrode on 40Hz oscillatory power on the whole brain level.

| Condition | Factor | d.f. | F value | p value | $\eta p^2$ |
| --- | --- | --- | --- | --- | --- |
| **Eyes-open** | stimulation | 4, 38638 | 236.19 | **<0.001** | 0.03 |
|  | time | 4, 38637 | 470.71 | **<0.001** | 0.05 |
|  | electrode | 60, 38637 | 8.94 | **<0.001** | 0.01 |
| | stimulation × time | 16, 38637 | 29.10 | **<0.001** | 0.01 |
| | stimulation × electrode | 240, 38637 | 2.19 | **<0.001** | 0.02 |
| | electrode × time | 240, 38637 | 1.06 | 0.24 | 0.00 |
| | stimulation × time × electrode | 960, 38637 | 0.50 | 1.00 | 0.01 |
| **Eyes-closed** | stimulation | 4, 38309 | 149.64 | **<0.001** | 0.02 |
|  | time | 4, 38308 | 735.70 | **<0.001** | 0.07 |
|  | electrode | 60, 38308 | 7.01 | **<0.001** | 0.01 |
| | stimulation × time | 16, 38308 | 17.01 | **<0.01** | 0.00 |
| | stimulation × electrode | 240, 38308 | 2.22 | **<0.001** | 0.01 |
| | electrode × time | 240, 38308 | 1.10 | 0.19 | 0.00 |
| | stimulation × time × electrode | 960, 38308 | 0.55 | 1.00 | 0.01 |

#### 3.3.1. Eyes open condition

Compared to the baseline, the X-tACS Peak protocol significantly increased 40Hz oscillatory power in the frontal and temporal regions of both hemispheres within the 0-30 minute window. The tACS protocol enhanced 40Hz power in the frontal, parietal, and occipital regions, while the iTBS protocol increased 40Hz oscillatory power in the right frontal and right parietal cortices. No significant effects were observed for the X-tACS Trough and sham stimulation protocols. In the 30-60 minutes time window, the X-tACS Peak and X-tACS Trough protocol significantly enhanced 40Hz oscillations, particularly in the frontal and temporal regions. The tACS protocol continued to demonstrate a significant increase in the target frequency power, particularly in the frontal and occipital regions. No significant changes were observed for the iTBS and sham stimulation protocols during this period.

During the 60-90 minutes interval, the X-tACS Peak protocol significantly increased 40Hz oscillatory power across the frontal, temporal, parietal, and occipital cortices. The significant effects from the previous time window for the X-tACS Trough intervention continued. The tACS intervention exhibited intensified effects, in 40Hz power across the same regions. For the iTBS protocol, a significant increase in 40Hz power was observed around the stimulation sides, extending to the right parietal cortex. No significant effects were noted for sham stimulation. In the 90-120 minutes post-intervention, significant effects from the X-tACS Peak, X-tACS Trough, and tACS protocols were maintained. The X-tACS Trough protocol also showed an additional significant increase in 40Hz oscillatory power in the left parietal cortex. The effects of the iTBS protocol intensified, revealing new significant increases in the occipital cortex, while sham stimulation showed no significant effects (Figure 3a).

**Figure 3:**
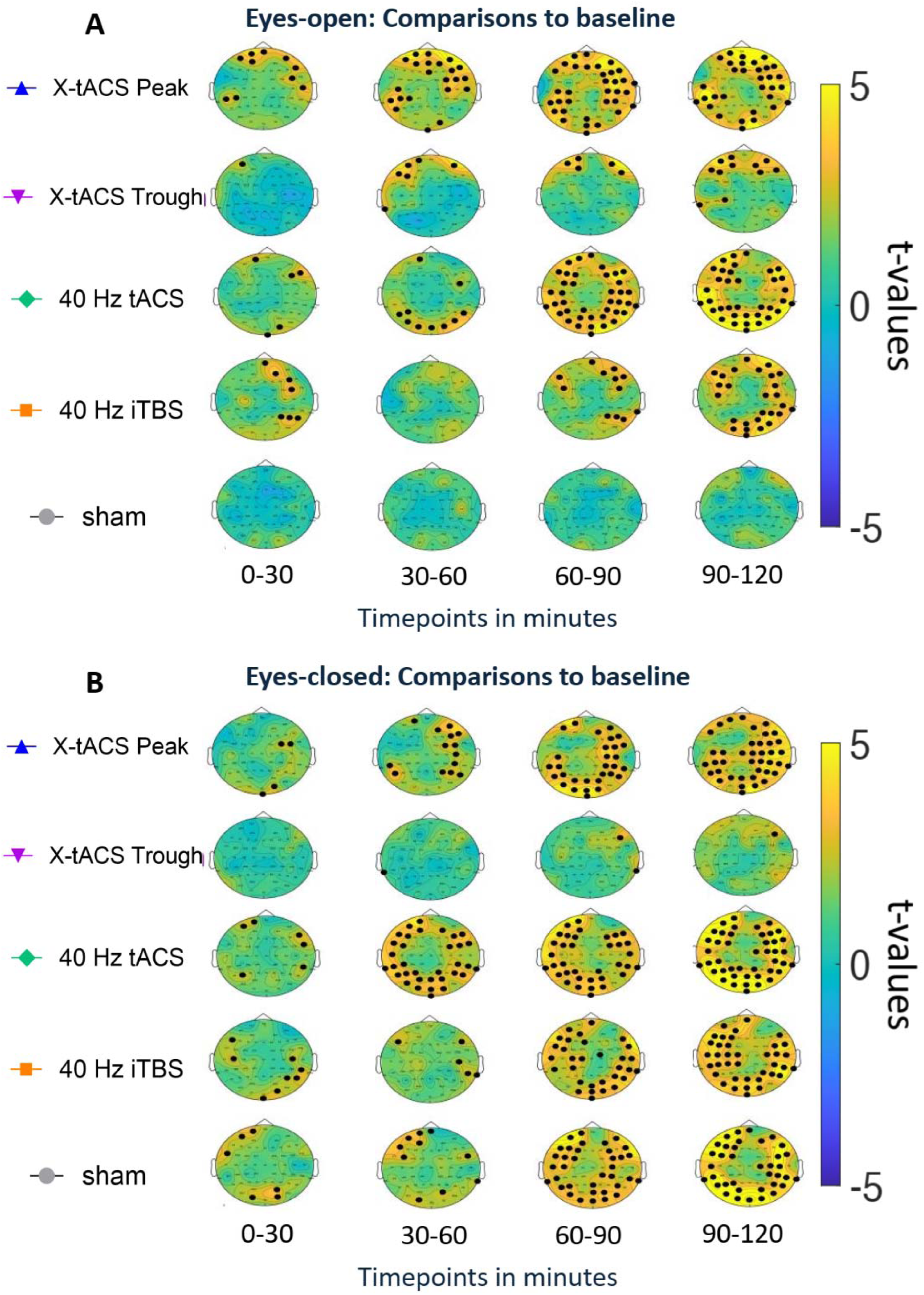
**A,** Whole brain changes in 40Hz oscillatory power are shown at four-time windows following the intervention (0-30, 30-60, 60-90, and 90-120 minutes) relative to baseline during **eyes-open** recordings. T-values from post hoc tests are shown on topological graphs. Positive t-values indicate an increase in oscillatory power, while negative values indicate a decrease. Black dots indicate a significant difference from baseline (p<0.05, FDR-corrected). **B,** Whole brain changes in 40Hz oscillatory power across different time windows (0-30, 30-60, 60-90, and 90-120 minutes) are compared to the baseline for **eyes-closed** recordings. The t-values from post hoc tests are plotted on topographical graphs, with positive t-values indicating an increase and negative values indicating a decrease in oscillatory power. Black dots represent significant differences from the baseline (p<0.05, FDR-corrected).

Compared to the sham condition, the X-tACS Peak protocol enhanced 40Hz oscillatory power in the frontal and temporal cortices of both hemispheres within 0-30 minutes post-intervention while the iTBS protocol effects on 40Hz oscillatory power included frontal and parietal cortices. For tACS, a significant 40Hz power increase was observed in only two frontal electrodes, while the X-tACS Trough protocol showed a significant decrease of respective oscillations in one right parietal electrode. The effects from the X-tACS Peak protocol were sustained in the 30-60 minute window, and a significant increase in 40Hz power was noted for the X-tACS Trough protocol in the frontal cortex, with no significant effects for tACS and iTBS. In the 60-90 minute period, the significant effects of the X-tACS Peak protocol intensified, with additional increases in the bihemispheric parietal and occipital cortices. The X-tACS Trough effects were sustained, and tACS also showed a significant increase in 40Hz power across the frontal, temporal, parietal, and occipital cortices. The iTBS protocol revealed a focal significant increase in oscillatory power around the stimulation sites and in the right parietal cortex. In the 90-120 minute window, all active stimulation protocols maintained and intensified their effects, with an additional significant increase in 40Hz power in the left parietal cortex for the X-tACS Trough protocol (Figure 4a).

**Figure 4:**
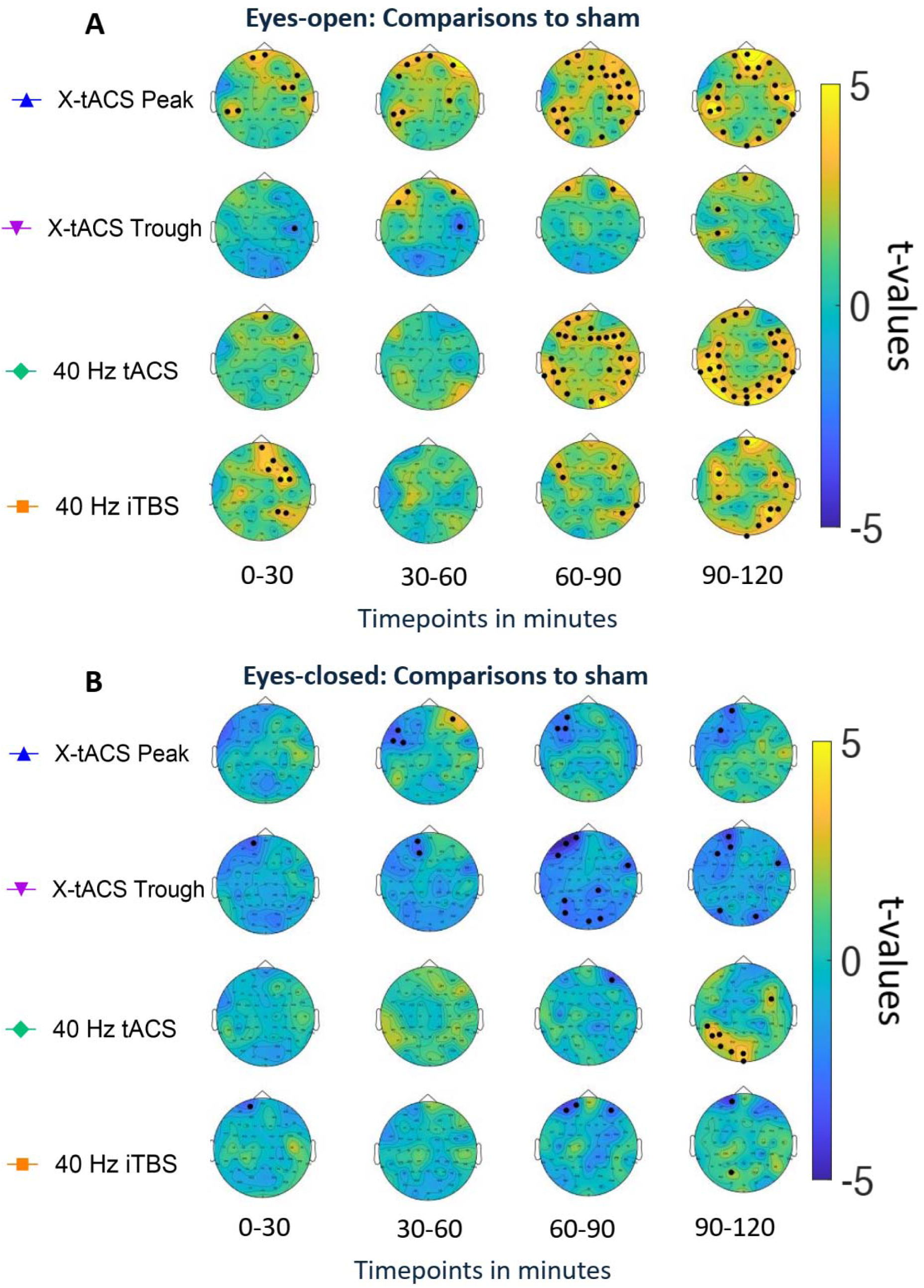
**A,** Whole brain changes of 40Hz oscillatory power for the different time windows (0-30 minutes, 30-60 minutes, 60-90 minutes, and 90-120 minutes) are presented compared to the sham stimulation for the **eyes open** recordings. t-values obtained from the post hoc tests are plotted onto the topological graphs. Positive t-values indicate an increase in oscillatory power, whereas negative values indicate a decrease in oscillatory power. Black dots indicate a significant difference compared to the sham condition (p<0.05, FDR-corrected). **B**) Whole brain changes of 40Hz oscillatory power for the different time windows (0-30 minutes, 30-60 minutes, 60-90 minutes, and 90-120 minutes) are presented compared to the sham stimulation for the **eyes closed** recordings. t-values obtained from the post hoc tests are plotted onto the topological graphs. Positive t-values indicate an increase in oscillatory power, whereas negative t-values indicate a decrease in oscillatory power. Black dots indicate a significant difference compared to the sham condition (p<0.05, FDR-corrected).

#### 3.3.2. Eyes closed condition

The X-tACS Peak significantly increased 40Hz power in the right frontal and occipital cortices within the first 30 minutes post-intervention, while tACS had similar effects in bilateral frontal and parietal cortices. iTBS increased 40Hz power in the left frontal and right parieto-occipital cortices. Sham stimulation increased 40Hz power in the left frontal and occipital regions, while X-tACS Trough showed no effect. Between 30-60 minutes, X-tACS Peak significantly enhanced 40Hz power in the right frontal, temporal, and bilateral parietal cortices, and tACS increased it across bilateral frontal, temporal, parietal, and occipital cortices. iTBS moderately increased 40Hz power in the frontal and right parietal regions. Sham stimulation also raised 40Hz power in the bilateral frontal, parietal, and occipital cortices; X-tACS Trough had no effect. From 60-90 minutes, the effects of X-tACS Peak and tACS persisted and intensified, particularly in the temporoparietal and occipital cortices for X-tACS Peak. iTBS showed stronger effects on 40Hz power in the frontal, temporal, parietal, and occipital regions, as did sham stimulation. X-tACS Trough still showed no effect. These effects persisted and intensified from 90-120 minutes (see Figure 3b).

There were no significant effects of stimulation protocols compared to the sham condition in the first 30 minutes post-intervention. In the 30-60 minute window, both the X-tACS Peak and X-tACS Trough protocols showed a significant decrease in 40Hz power in the left frontal region, with no other significant effects detected. This decrease persisted in the 60-90 minute period for both protocols, alongside significant decreases in the right temporal, left parietal, and occipital regions. The iTBS protocol also led to a significant decrease in frontal regions, with no other significant effects found. In the final 90-120 minute window, previous effects persisted, and a significant increase in 40Hz power was observed in the left parieto-occipital region for the tACS protocol, with no other significant effects noted (Figure 4b).

## 4 Discussion

This study investigated the impact of five NIBS protocols, developed via a combination of tACS and iTBS, on inducing and stabilizing 40Hz gamma oscillations in healthy individuals. Specifically, we investigated the impact of 40-Hz tACS, 40-Hz iTBS, phase-locked 40 Hz iTBS to 40 Hz tACS peak sine wave, phase-locked 40 Hz iTBS to 40 HZ tACS trough sine wave, and a sham iTBS-tACS over the left and right dorsolateral prefrontal cortex. In addition to examining the local effects of the stimulation in the frontal lobe, we assessed the global impact of NIBS on 40Hz gamma power across the entire brain. In what follows, we discuss major physiological findings.

### 4.1. 40 Hz entrainment under NIBS interventions and induced gamma oscillations

Our major finding is that combined protocol (40Hz iTBS phase-locked to tACS) most effectively induced gamma activity in the EO recording, followed by 40Hz tACS and 40Hz iTBS alone. In the EC condition, iTBS protocols significantly decreased 40Hz power compared to sham, while tACS alone most effectively boosted 40Hz power between 30-90 minutes. The gamma-enhancing effects of tACS applied at the peak of oscillations partly replicate previous findings showing enhanced alpha and theta power in the EO condition using the same intervention [29], [30], although the opposite pattern was observed for the delta band with tACS applied at the trough [31].

A possible explanation for the primary finding is that iTBS applied at the sine wave may have induced in-phase 40Hz neuronal activity, facilitating tACS entrainment. Previous research has demonstrated that repetitive rTMS can transiently alter oscillations based on pulse frequency, causing a “phase reset” [7], [44] followed by short-lasting synchronization at the driving frequency [8]. Thus, iTBS-induced 40Hz activity may enhance alignment between endogenous and external oscillations, strengthening tACS entrainment. Additionally, some studies emphasize the importance of ongoing oscillatory activity during tACS, which iTBS can modulate. Kemmerer et al. [45] showed that tACS at the individual alpha peak significantly modulated oscillatory power in that frequency, whereas +/− 2Hz or sham stimulation was less effective, highlighting tACS’s frequency specificity. Similarly, frontoparietal tACS aligned with the individual theta peak, compared to non-individualized 4Hz tACS or sham, led to stronger oscillatory power and network synchrony [46]. Consistent with this, the strongest modulation occurred at the 40Hz target frequency, confirming tACS’s capacity to selectively entrain endogenous oscillations in a frequency-dependent manner. While oscillatory effects were also present in other bands like theta and alpha (Figure S4), they were less pronounced.

Another explanation concerns increased efficacy of iTBS to induce action potentials at the cortical level when the TMS pulses are applied to the peak of the tACS sine wave. TACS aligns ongoing natural oscillatory activity with its exogenous current [10]. Non-human primate studies, along with computational models, show that as oscillatory activity becomes aligned, the underlying neurons exhibit stronger depolarization during the positive phase (peak) of the applied current, while a hyperpolarized state is observed during the negative phase (trough) of the current [36], [37], [38]. Applying iTBS at the peak, when the neuronal membrane is depolarized and closer to the firing threshold, may facilitate action potential generation. Conversely, iTBS at the trough, during hyperpolarization, would likely be less effective. Consistent with this, Wischnewski et al. [47] showed that cortical excitability depends on the phase of ongoing oscillations, with TMS pulses delivered during the peak or falling phase of the beta rhythm resulting in increased MEP amplitudes. This highlights the enhanced efficacy of TMS pulses during the depolarized phase and explains the stronger effects observed with the X-tACS Peak protocol.

Although the most pronounced increase in gamma oscillatory power occurred in the iTBS phase-locked to the peak of the tACS wave during eyes eyes-open state, the specific mechanisms driving the differential effects of these interventional protocols require further investigation. A difference of the results of this study compared to our previous works [29], [30] is that the stimulation effects did not transfer to the EC recording condition, while the same effect was seen for both EO and EC conditions in those studies. One possible reason for the greater intervention effects in the EO condition in the present study is the stronger high-frequency brain activity, including gamma rhythms, present when the eyes are open, compared to the general shift to lower frequencies, such as alpha rhythms, when the eyes are closed [48]. The stronger effects observed in the EO condition may be due to the higher frequency oscillatory activity during the baseline state at the time of stimulation, which occurred during rest with the eyes open. Prior research indicates that baseline oscillatory activity in the target frequency is critical for the entrainment effects of tACS [47]. Similarly, in rTMS, the baseline brain state during stimulation influences oscillatory changes [49], which may explain the enhanced effects in the EO condition.

### 4.2. Gamma oscillations under other interventions and brain states

The results of resting EEG analyses specifically during the eyes closed condition and overall (eyes open + eyes closed) strongly show that the 40 Hz tACS alone induced 40 Hz gamma power most effectivity and consistently up to 2 hours post-intervention. This may be because the eyes-closed state provides an arousal baseline, while the eyes-open condition provides an activation baseline [48]. The superior effectiveness of 40 Hz tACS alone in inducing gamma power during eyes closed indicates tACS’s ability to linearly increase gamma power over time, when the brain is not further activated by visual input. This suggest that tACS might be the most effective protocol, when brain activity and endogenous power oscillations is lower [50], tACS has been the most effective Conversely, increased arousal and visual engagement in the eyes-open state may amplify the effectiveness of all interventions, diminishing the relative superiority of tACS alone seen in the eyes-closed condition. The similar effects of tACS and the Peak protocol within the first 30 minutes of the eyes-open condition, with iTBS showing the largest effect, further suggest that the activation baseline allows multiple interventions to effectively modulate gamma power. The strong acute effect of 40 Hz iTBS and tACS alone (in the first 30 minutes) is another worth noting observation.

### 4.3. Limitations, future directions, and therapeutic potential

The present study has several limitations. Firstly, it served as a proof-of-concept, examining the effects of the new stimulation protocols on oscillatory aftereffects and safety in a healthy population. Although no adverse events were reported, further research is needed to explore the stimulation effects across different healthy age groups and in clinical populations, such as those with mild cognitive impairment or Alzheimer’s disease, to enhance therapeutic outcomes and explore whether clinical translation is feasible or not. Additionally, we cannot discount the possibility that visual phenomena from the tACS intervention influenced the observed 40Hz oscillatory changes in the frontal region of interest. Future studies should address these visual effects, especially with higher stimulation intensities, which may exacerbate them. Moreover, effective blinding techniques should be implemented to reduce the confounding effects of phosphenes during tACS application. Lastly, although our sample size aligns with previous studies and power analyses, increasing the sample size in future research is recommended to enhance statistical power, particularly in studies involving clinical populations.

The primary objective of the study was to evaluate the potential of various 40 Hz entraining NIBS interventions in inducing gamma oscillations, which are typically reduced in AD. While future studies should explore this in actual AD patients, the current findings can be discussed in terms of clinical efficacy and feasibility. The combined 40 Hz iTBS-tACS (phase-locked at the peak of the tACS wave) and 40 Hz tACS alone demonstrated stronger and longer-lasting induced gamma oscillations, making them more promising for therapeutic applications in AD. Regarding aftereffects, these protocols sustained gamma oscillations for up to two hours and exhibited linearly increasing effects during eyes-open conditions (combined protocol) and eyes-closed conditions (tACS alone). Nonetheless, a strong acute effect of the iTBS-alone intervention on gamma oscillations was also observed (during eyes-open and overall resting EEG states). However, the lack of prolonged aftereffects suggests that iTBS alone may not be as effective as the combined 40 Hz iTBS-tACS or 40 Hz tACS alone. Regarding feasibility, the combined protocols require a complex setup involving two TMS machines, one tES device, and a neuronavigator. While this setup may be practical in a laboratory setting, it might not be feasible or affordable for some clinics. Consequently, the use of 40 Hz tACS alone, which can induce gamma oscillations for up to two hours, appears to be a more practical option. Alternatively, a modified approach to the combined protocol, such as stimulating only one hemisphere, should be explored in future studies to assess its comparative efficacy.

### 4.4. Conclusion

In conclusion, all protocols increased 40Hz gamma oscillations during eyes-open recordings. The most effective protocol was the X-tACS Peak protocol, where the iTBS pulses were phase-locked to the peak of the tACS-generated sine wave. This finding is in accordance with findings from previous research with similar stimulation approaches. During eyes closed and overall state, the 40 Hz tACS alone was the most effective and consistent protocol in inducing gamma oscillation. Future studies should focus on the underlying mechanisms of the aftereffects produced by the stimulation protocols and implications for the transfer to behavioral modulations.

## Consent Statement

All subjects provided written informed consent

## CrediT authorship contribution statement

**Benedikt Glinski:** Data curation, Formal analysis, Investigation, Methodology, Writing – original draft, Visualization. **Mohammed Ali Salehinejad:** Data curation, Visualization, Formal analysis, Writing – original draft, Methodology, Writing – review & editing. **Kuri Takahashi:** Data curation, Writing – review & editing. **Asif Jamil**: Conceptualization, Writing – review & editing. **Fatemeh Yavari:** Methodology, Writing – review & editing. **Min-Fang Kuo:** Conceptualization, Methodology, Supervision, Validation, Resources, Funding acquisition, Writing – review & editing. **Michael A. Nitsche:** Conceptualization, Supervision, Methodology, Resources, Funding acquisition, Writing– review & editing.

## Declaration of competing interest

The authors declare the following financial interests/personal relationships which may be considered as potential competing interests: MAN is a member of the Scientific Advisory Boards of Neuroelectrics and Precisis. None of the remaining authors have potential conflicts of interest to be disclosed.

## Funding

This project has received funding from the Horizon 2020 research and innovation program of the European Union under grant agreement No 101017716 (Neurotwin)

### Acknowledgment

We thank Daniel Strobel, Tobias Blanke, and Nina Abich for their technical support.

## Notes

### Summary of Updates

Revised version after first round of peer review

