## Supplementary material for "Phase-synchronized 40Hz tACS and iTBS effects on gamma oscillations"

**Supplementary materials**

**Methods**

***Intervention***

*Repetitive transcranial magnetic stimulation protocol*

Due to safety concerns, it is advised to apply classical rTMS with frequencies no higher than 25Hz [2]. Thus, a different protocol was used to stimulate in the gamma range (40Hz), specifically intermittent theta burst stimulation (iTBS), which is regarded as safe for high-frequency neuromodulation. Compared to classical rTMS, where pulses are applied repeatedly at a fixed frequency, iTBS consists of bursts of pulse triplets. In the classical protocol, the bursts are applied for two seconds in 5Hz intervals followed by an eight-second break, before the next bursts are applied. This pattern is repeated for 190 seconds resulting in 600 pulses per protocol. The high-frequency component of this protocol is the intra-burst interval of the triplets which in the classical protocol are applied at 50Hz. To match our target frequency of 40Hz, we changed the stimulation frequency from 50Hz to 40Hz. The rest of the protocol was kept the same as the classical one. The modified iTBS protocol was applied at 80% of the individual subjects’ active motor threshold (AMT) four times within a 20-minute time frame starting at 0 minutes, 5 minutes, 10 minutes, and 15 minutes from the beginning of the intervention resulting in 2400 delivered pulses for the whole intervention.

The stimulation intensity for iTBS was defined by first applying single-pulse biphasic TMS pulses over the left primary motor cortex representation of the abductor digit minimi muscle (ADM) to identify the motor cortical hotspot. The figure of eight coil (PMD coil type, Mag&More GmbH, Munich, Germany, diameter of one winding 70mm, peak magnetic field 2T) connected to a PowerMag stimulator (Mag&More GmbH, Munich, Germany) was held 45° to the midline over the target regions. First, the hotspot (the coil position over the primary motor area that produces the largest MEP in the right ADM) was identified with TMS in line with previous works [3,4]. All EMG signals were amplified and filtered (1000; 3Hz-3kHz) using the D440-2 (Digitimer, Welwyn Garden City, UK) and were digitized (sampling rate 5kHz) with a micro 1401 AD converter (Cambridge Electronic Design, Cambridge, UK), controlled by Signal Software (Cambridge Electronic Design, v. 2.13). Afterwards, the AMT was identified by applying single TMS pulses over the predefined motor hotspot during moderate tonic contraction of the ADM (defined as 20% of the EMG amplitude produced by maximal muscle contraction). The TMS pulse intensity was adjusted up ti the lowest intensity at which at least three out of six pulses elicited an MEP amplitude of ~ 200-300mV. 80% of the individual AMT was used as intensity for the experimental theta burst application [5].

*Transcranial alternating current stimulation*

tACS was delivered utilizing a Starstim 8-channel constant current, battery-powered electric stimulator (Neuroelectrics, Barcelona, Spain). Circular carbon rubber electrodes (2cm radius, 12,57 cm^2^) were used for stimulation. A topical anesthetic cream (EMLA®, Dublin, Irland) was applied to the skin over the target region to reduce stimulation-induced phenomena (e.g., tingling) and discomfort. Ten20 paste (Weaver and Company, USA) was applied to the bottom of the electrodes before placing the electrodes on the head to ensure good current conductivity. During tACS, current was delivered with a 1mA Peak-to-peak amplitude with a 15-second ramp up at the beginning of the stimulation and a 15-second ramp down at the end of the stimulation. Sham tACS was applied by only ramping the current for 15 seconds up and 15 seconds down at the beginning and end of the stimulation respectively.

*Combined stimulation protocols*

A customized circuit was developed in-house to combine tACS and the modified iTBS protocol by phase-locking iTBS pulses to the tACS waveform. iTBS pulses were delivered using two figure-of-eight-shaped TMS coils. For sham iTBS, two different sham TMS coils (double 70mm pCool-SHAM coil, Mag&More GmbH, Munich, Germany) generated audible clicks at the same intensity as an active coil without inducing an electromagnetic pulse. To minimize the delay between iTBS pulses from the two coils, the TMS devices were connected in series, with TTL signals used for triggering. An oscilloscope confirmed near-zero latency between the pulses.

The Starstim device used for tACS was modified for external stimulation pattern configuration. Two tACS channels were programmed to deliver square pulses at specific intervals relative to the tACS sine waves, while four additional channels provided sinusoidal alternating currents to the dorsolateral prefrontal cortex bilaterally at electrode positions F3 and F4, with return electrodes at the ipsilateral mastoids (TP9 and TP10) using the international 10-20 EEG system. An internally developed circuit with an optocoupler transmitted square pulses (0.1 mA peak, 0.5% duty cycle) from the electric stimulator to trigger the iTBS pulses, ensuring isolation from the main power. The current-to-voltage conversion produced a 5V signal for TTL communication with the TMS machine's TRIGGER-IN port. Synchronization of pulses and tACS sine waves was confirmed with an oscilloscope. During stimulation, the TMS coils were positioned over the F3 and F4 electrodes at a 45-degree angle to the midline, with coil positions marked using a commercial neuronavigation system (LOCALITE GmbH, Germany). The coil angle and deviation from marked positions were monitored throughout the stimulation.

***Data analysis***

*Physiological data analysis*

Preprocessed EEG power data were exported from Brainvision Analyzer for the target frequency of 40Hz, low Gamma (30-45Hz), Beta (14-30Hz), Alpha (8-13Hz), Theta (4-7Hz), and Delta (1-3Hz). Two subjects were removed from the final analysis due to strong physiological artifacts. Thus, the final EEG power analysis was performed on 28 subjects. The raw data were first baseline-standardized by dividing the post-stimulation power by the baseline power for each electrode at each measurement timepoint (note that absolute 40Hz baseline power was assessed for differences between conditions before standardization (Figure S3). Subsequently, the baseline-standardized data were collapsed by averaging the data into time windows of 0-30 minutes (T1-T4), 30-60 minutes (T5-T7), 60-90 minutes (T8-T10), and 90-120 minutes (T11-T13). Data points emerging from non-neurogenic sources were removed from the baseline-standardized data using the ROUT algorithm implemented in Prism [6] with Q set to 1%. This led to the removal of 4.68% of the whole data (for an example of pre-post data removal distributions and model fit see Figure S1a,b).

The baseline-standardized and epoched data were analyzed using linear mixed-effect models, fitted with the lmer function from the lme4 R package. The fixed effects included the stimulation protocol, EEG measurement time window, EEG channel, and their interactions. A by-participant random intercept term was included to account for the non-independence of observations within subjects. The significance levels for the model's fixed effects were computed using Satterthwaite's approximation for degrees of freedom and assessed using F-tests from the ANOVA function in the lmerTest R package. The critical significance level (alpha error) for all tests was set to 0.05.

In case of significant main or interaction effects, post-hoc pairwise comparisons were conducted using the estimated marginal means obtained from the model via the emmeans R package. These means accounted for both the random effects and main factors of the model. All post-hoc tests were corrected utilizing the false-discovery rate (FDR; [7]).

*Blinding and side effects analysis*

To explore blinding efficacy, participants were asked to identify whether they received real or sham stimulation after each condition. Fisher's Exact Test analyzed their ability to distinguish real stimulation types (X-tACS Peak, X-tACS Trough, tACS, iTBS) from the sham condition. We also evaluated side effects—such as tingling, burning, pain, skin redness, and visual phenomena—using a questionnaire where participants rated the intensity of each symptom on a Likert scale from 0 (absent) to 5 (very strong). The results were analyzed using a one-way repeated measures ANOVA for each phenomenon. In case of significant main effects, post hoc t-test were used to further investigate the main effects.

**Results**

***Effects of stimulation on other frequency bands***

Next, we investigated the effects of the stimulation on other frequency bands outside the target frequency in the frontal ROI. Linear mixed-effects models were defined as described before. For the delta (1-3Hz) band, significant main effects were found in the EO condition for stimulation (F_4,7228_ = 4.66, p<0.001), time (F_1,7228_= 99.74, p<0.001) a significant stimulation × time interaction (F_4,7227_= 4.66, p<0.001) was also revealed. In the EC condition main effects seen for time (F_4,7317_ = 31.21, p < 0.001) electrode (F_10,7317_ = 2.19, p < 0.01) and an interaction effect of time × electrode (F_10,7317_ = 2.19, p < 0.05) was found.

Compared to the baseline, all protocols except for sham showed an increase in delta power (all p < 0.05). In the EO condition, post hoc pairwise comparisons revealed that X-tACS Peak, X-tACS Trough, and tACS significantly elevated delta power compared to both sham and the iTBS protocol (all p < 0.05). Notably, iTBS did not differ significantly from sham (t = 2.02; p = 0.053). Additionally, X-tACS Trough produced a greater increase in delta power than the tACS protocol (t = 3.14; p = 0.002). In the EC condition all protocols with the exception of iTBS (t = 1.15; p = 0.24) increased delta power relative to baseline (all p < 0.05).

Compared to sham only tACS led to an increase of delta power (t = 2.50; p = 0.03).

In the theta (4-7Hz) band, significant main effects were seen for stimulation (EO: F_4,7326_= 6.44, p<0.001; EC: F_4,7351_=2.95, p<0.001), time (EO: F_1,7326_= 334.34, p<0.001; EC: F_4,7351_=258.03, p<0.001), and electrode (EO: F_10,7326_ =3.78, p < 0.001; EC: F_10,7351_=4.25, p<0.001) a significant stimulation × time (EO: F_4,7326_= 6.44, p<0.001; EC F_4,7351_=2.95, p<0.001), and a significant electrode × time interaction (EO: F_10,7326_= 3.78, p<0.001; EC F_10,7351_=4.25, p<0.001) was found. In the EO condition, all interventions increased theta power relative to baseline(all p < 0.001). Compared to sham all interventions significantly increased theta power (all p < 0.05). X-tACS Trough demonstrated the strongest effect, surpassing all other protocols (all p < 0.01). No significant differences were observed among the other protocols. In the EC condition, all protocols, including sham, led to a significant increase in theta power relative to baseline (all p < 0.001). Compared to sham, X-tACS Peak, X-tACS Trough, and tACS significantly elevated theta power (all p < 0.05).

For the alpha (8-13Hz) band, significant main effects were found in the EO and EC conditions for stimulation (EO: F_4,7305_ = 13.31, p<0.001; EC: F_4,7275_ = 4.54, p<0.001), time (EO: F_1,7305_= 1153.33, p<0.001; EC: F_1,7275_= 339.45, p<0.001) and the interaction of stimulation × time was significant (EO: F_4,7305_= 13.31, p<0.001; EC: F_4,7275_= 99.74, p<0.001). In addition, a significant main effect was found for electrode (F_10,7305_ = 2.65, p<0.001) and the interaction of electrode × time was significant (F_10,7305_ = 2.65, p<0.001) in the EO condition. In the EO condition, all protocols significantly increased alpha power relative to baseline (all p < 0.001). Compared to sham all interventions significantly elevated alpha power (all p < 0.001). Notably, the X-tACS Trough protocol showed the greatest increase in alpha power among all interventions (all p < 0.01). Furthermore, iTBS produced a significantly stronger effect than both X-tACS Peak and tACS (all p < 0.001). No significant difference was observed between the tACS and X-tACS Peak protocols (t = 0.41; p = 0.68). In the EC condition all protocols increased alpha power relative to baseline (all p < 0.001). Compared to sham, X-tACS Peak, X-tACS Trough, and iTBS significantly reduced alpha oscillations (p`s < 0.05). No significant difference was observed between tACS and sham (t = 0.42; p = 0.67). The X-tACS Trough protocol resulted in the most substantial reduction in alpha power, significantly surpassing all other protocols (all p < 0.001). Additionally, X-tACS Peak induced a greater decrease than the iTBS protocol (t = 2.17; p = 0.03).

In the beta band (14-30Hz) significant main effects were found for stimulation (EO: F_4,7265_= 6.42, p<0.001; EC: F_4,7335_=2.86, p<0.05) and time (EO: F_1,7265_= 346.58, p<0.001; EC: F_4,7335_=507.78, p<0.001). Furthermore, a significant stimulation × time interaction (EO: F_4,7265_= 6.42, p<0.001; EC F_4,7335_=2.86, p<0.05) was found. No additional significant main or interaction effects were found. Post hoc pairwise comparisons in the EO condition indicated that all protocols significantly enhanced beta power relative to baseline (all p < 0.001). Compared to sham, all protocols significantly increased beta power (all p < 0.001). X-tACS Peak and X-tACS Trough resulted in smaller beta power increases than iTBS (all p < 0.01). In the EC condition, all protocols led to a significant increase of beta power relative to baseline (all p < 0.01). Compared to sham, X-tACS Peak and iTBS significantly decreased beta power (all p < 0.001). X-tACS Trough and tACS showed no significant effect relative to sham. X-tACS Peak resulted in a greater decrease in beta power compared to all other protocols, including iTBS (all p < 0.01). Additionally, X-tACS Trough significantly reduced beta power compared to tACS (t = 3.10; p < 0.01). No further effects were observed.

With respect to low gamma (30-45Hz) power, LMM analyses showed significant main effects of stimulation in the EO and EC conditions (EO: F_4,7217_ = 9.52, p<0.001; EC: F_4,7276_ = 4.66, p<0.001), time (EO: F_1,7217_= 236.78, p<0.001; EC: F_1,7276_= 316.38, p<0.001) and a significant interaction of stimulation × time (EO: F_4,7217_= 9.52, p<0.001; EC: F_4,7276_= 4.66, p<0.001). In addition, a significant main effect was found for electrode (EO: F_10,7217_ = 2.09, p<0.05) and the electrode × time interaction (F_10,7217_ = 2.09, p<0.05) in the EO condition was significant. In the EO condition, all protocols significantky increased gamma power relative to baseline (all p < 0.001). Compared to sham, all protocols significantly increased low gamma power (all p < 0.001). The X-tACS Peak protocol produced the most substantial increase relative to all other protocols (all p < 0.01). Additionally, iTBS led to a stronger increase than both tACS and X-tACS Trough (all p < 0.001), with no significant difference observed between the latter two (t = 0.31; p = 0.76). In the EC condition, all protocols significantly increased gamma power relative to baseline (all p < 0.001). Compared to sham, pairwise comparisons revealed that X-tACS Peak and X-tACS Trough significantly decreased gamma power (all p < 0.001). No significant differences were observed between tACS and sham (t = 1.98; p = 0.06), or between X-tACS Peak and X-tACS Trough (t = 1.21; p = 0.25) . However, iTBS significantly increased gamma power more than tACS (t = 2.17; p = 0.04).

**Figures**

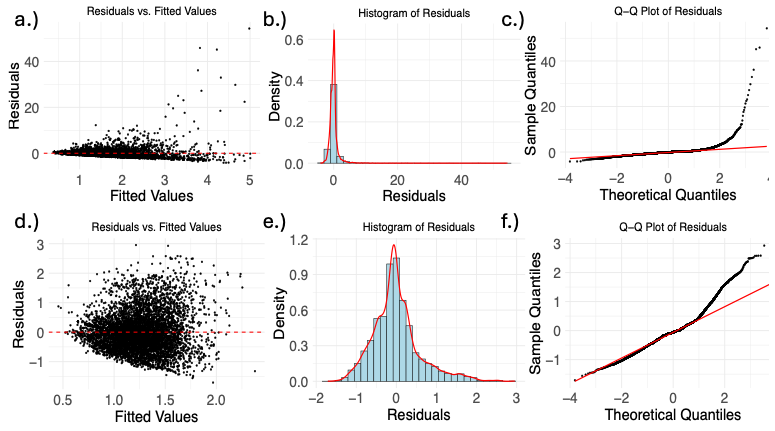

Figure S1: Different visualization of model fit before and after removing outliers with the ROUT algorithm based on an example from the frontal ROI 40Hz power data. **a/d.)** These scatterplots visualize the residuals of the linear mixed model (LMM) against the fitted values. Each point represents an observation, with the y-axis showing the residual (i.e., the difference between observed and predicted values). The red horizontal line at y axis (0) indicates the expected residual mean under a well-fitted model. Ideally, residuals should be **randomly scattered** around this residual mean of 0 without patterns to indicate a good model fit. **b/e.)** These histograms displays the distribution of residuals linear mixed model (LMM). The x-axis represents the residual values, while the y-axis shows the density of these residuals. The histogram bars (filled in blue) illustrate the frequency of residuals across different values, while the overlaid red density curve provides a smooth approximation of the residual distribution. A symmetric, bell-shaped distribution centered around zero suggests that the residuals follow a normal distribution. Deviations from normality, e.g., heavy tails (as in figure 1b.), are an indicator of bad model fit . **c/f.)** These Q-Q (quantile-quantile) plots display are another representation of the model fit. The x-axis represents the theoretical quantiles expected under normality, while the y-axis displays the sample quantiles of the actual LMM. If the residuals follow a normal distribution, they should closely align with the red diagonal line. Strong and systematic deviations from this line, e.g., heavy tails, suggest a non-normal distribution, which indicate bad model fit, outliers, or non-constant variance.

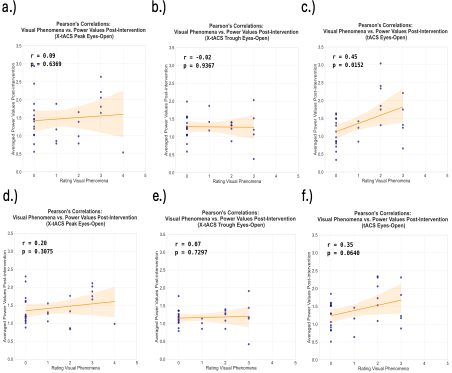

Figure S2a: Pearson correlations between visual phenomena and post intervention standardized power. The first panel shows the correlations for the different stimulation protocols on the frontal ROI, whereas the second panel shows the correlations on the whole brain level during Eyes-open recordings.

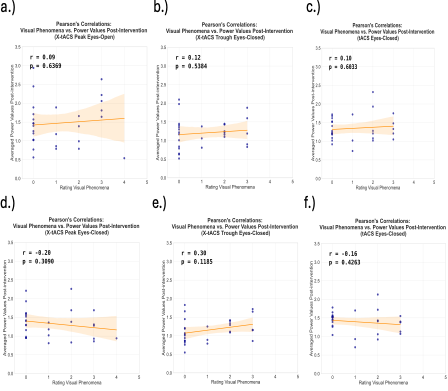

Figure S2b: Pearson correlations between visual phenomena and post intervention standardized power. The first panel shows the correlations for the different stimulation protocols on the frontal ROI, whereas the second panel shows the correlations on the whole brain level during Eyes-closed recordings.

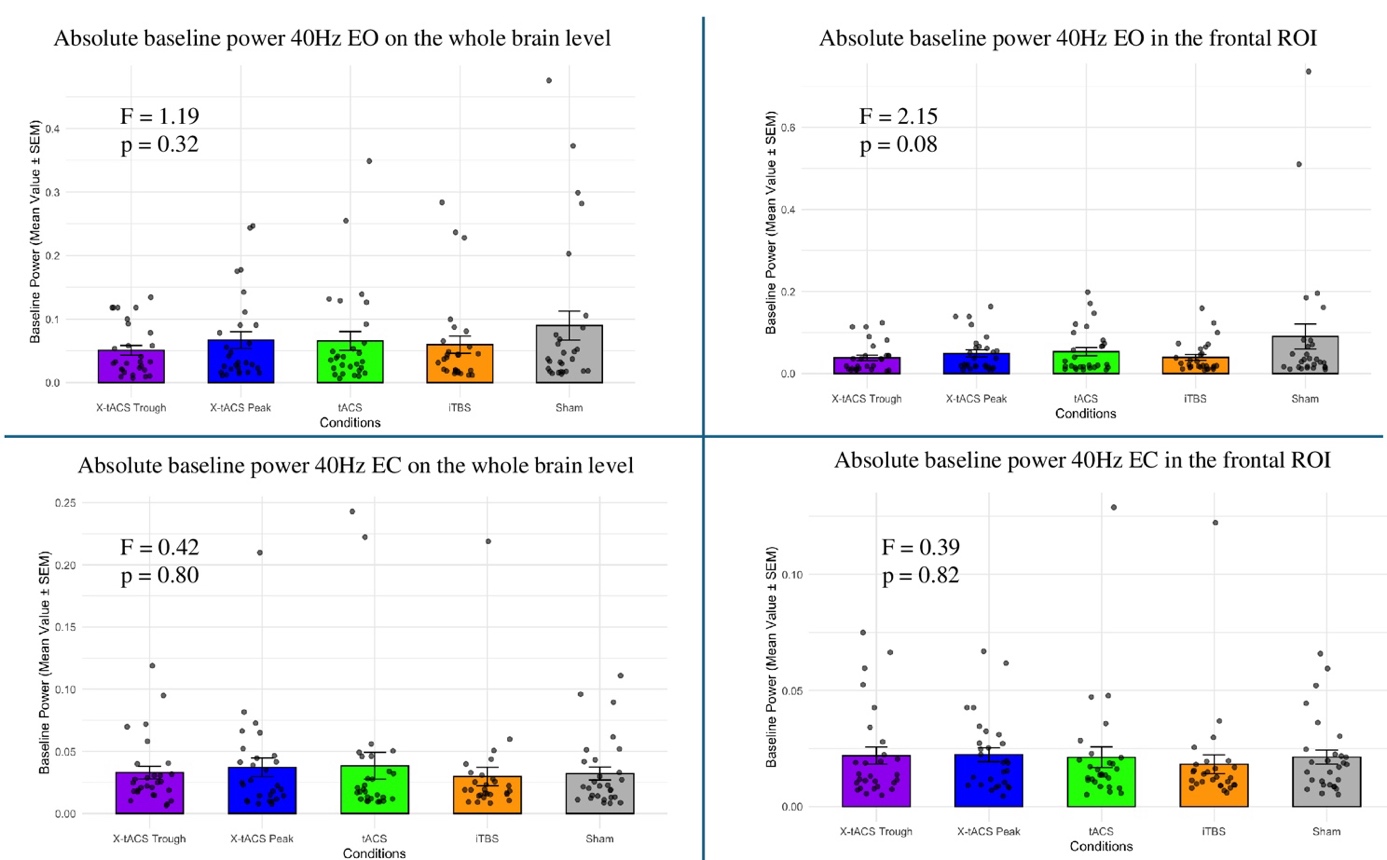

Figure S3: Absolute baseline power EO and EC for frontal ROI and whole brain level

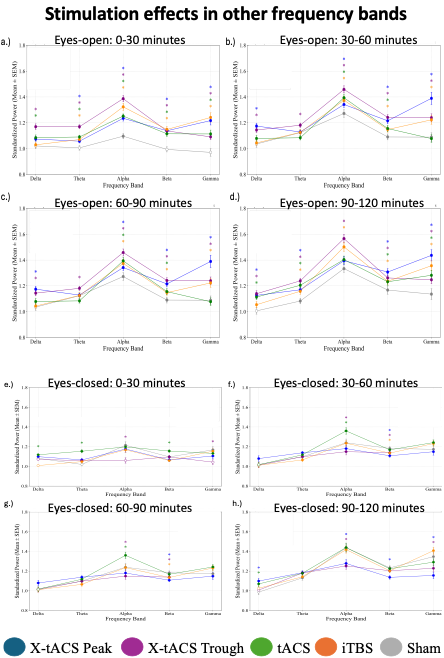

Figure S4. The effects of interventions on other frequency bands

**Tables**

**Table S1:** Fisher's exact Test to assess blinding efficacy

| **Stimulation conditions** | ***p*** |
| --- | --- |
| X-tACS Peak vs sham | 1.0000 |
| X-tACS Trough vs sham | 0.5047 |
| tACS vs sham | 0.7458 |
| iTBS vs sham | 1.0000 |

**Table S2.** Intensity ratings of the sensations per stimulation protocol in the Resting-State EEG Sessions. Data are presented as mean ± SD.

|  | |  | | **X-tACS Peak** | | **X-tACS Trough** | | **tACS** | | **iTBS** | | **Sham** |
| --- | --- | --- | --- | --- | --- | --- | --- | --- | --- | --- | --- | --- |
| **During stimulation** | Visual phenomena | | 1.10±1.31 | | 1.19±1.41 | | 1.07±1.21 | | 0.21±0.68 | | 0.18±0.47 | |
|  | Itching | | 0.11±0.41 | | 0.34±0.75 | | 0.60±1.13 | | 0.35±0.86 | | 0.21±0.62 | |
|  | Tingling | | 0.28±0.53 | | 0.23±0.71 | | 0.50±1.07 | | 0.17±0.66 | | 0.10±0.41 | |
|  | Burning | | 0.25±0.64 | | 0.03±0.19 | | 0.14±0.52 | | 0.17±0.54 | | 0±0 | |
|  | Pain | | 0.21±0.62 | | 0.57±1.17 | | 0.14±0.75 | | 0.35±0.78 | | 0.14±0.59 | |
| **After end of experiment** | Skin redness | | 0±0 | | 0±0 | | 0±0 | | 0±0 | | 0±0 | |
| **24 hours after stimulation** | Headache | | 0.11±0.41 | | 0.08±0.39 | | 0.03±0.19 | | 0.32±0.86 | | 0.21±0.68 | |
|  | Fatigue | | 0.57±0.92 | | 0.92±1.41 | | 0.82±1.33 | | 0.78±1.28 | | 0.71±1.27 | |
|  | Difficulty in concentration | | 0.17±0.55 | | 0.46±1.17 | | 0.28±0.65 | | 0.39±0.65 | | 0.17±0.66 | |
|  | Nervousness | | 0±0 | | 0.07±0.39 | | 0±0 | | 0±0 | | 0±0 | |
|  | Sleep problems | | 0±0 | | 0±0 | | 0±0 | | 0±0 | | 0±0 | |
|  | Others | | 0±0 | | 0±0 | | 0±0 | | 0±0 | | 0±0 | |

**Table S3**. One-way repeated-measures ANOVAs for adverse effects in the Resting-EEG sessions.

|  | ***Factor*** | ***d.f., error*** | ***F value*** | ***η^2^_p_*** | ***p value*** |
| --- | --- | --- | --- | --- | --- |
| **During stimulation** | Visual phenomena | 4,133 | 6.15 | 0.16 | **0.0001** |
|  | Itching | 4,133 | 1.55 | 0.04 | 0.19 |
|  | Tingling | 4,133 | 1.22 | 0.04 | 0.31 |
|  | Burning | 4,133 | 1.39 | 0.04 | 0.24 |
|  | Pain | 4,133 | 1.34 | 0.04 | 0.46 |
| **After end of experiment** | Skin redness | 4,133 | N/A | N/A | N/A |
|  | Headache | 4,133 | 1.17 | 0.03 | 0.33 |
|  | Fatigue | 4,133 | 0.30 | 0.01 | 0.88 |
|  | Difficulty in concentration | 4,133 | 0.74 | 0.02 | 0.57 |
|  | Nervousness | 4,133 | 1.08 | 0.03 | 0.37 |

df = degrees of freedom, η^2^p = partial eta squared. N/A = all responses equaled 0. No variance in the data.

3.2.1 *Resting EEG – Eyes closed vs eyes open*

**Table S4:** Pairwise comparisons between conditions 40Hz eyes-open

| **contrast** |  | **estimate** | **SE** | **df** | **t-value** | **p-value** | **d** |
| --- | --- | --- | --- | --- | --- | --- | --- |
| X-TACS TROUGH - SHAM | BL | 0,00 | 0,05 | 6978,00 | 0,00 | 1,0000 | 0,00 |
| X-TACS PEAK - SHAM | BL | 0,00 | 0,05 | 6978,00 | 0,00 | 1,0000 | 0,00 |
| X-TACS Peak - X-TACS TROUGH | BL | 0,00 | 0,05 | 6978,00 | 0,00 | 1,0000 | 0,00 |
| TACS - SHAM | BL | 0,00 | 0,05 | 6978,00 | 0,00 | 1,0000 | 0,00 |
| TACS - X-TACS TROUGH | BL | 0,00 | 0,05 | 6978,00 | 0,00 | 1,0000 | 0,00 |
| TACS - X-TACS PEAK | BL | 0,00 | 0,05 | 6978,00 | 0,00 | 1,0000 | 0,00 |
| ITBS - SHAM | BL | 0,00 | 0,05 | 6978,00 | 0,00 | 1,0000 | 0,00 |
| ITBS - X-TACS TROUGH | BL | 0,00 | 0,05 | 6978,00 | 0,00 | 1,0000 | 0,00 |
| ITBS - X-TACS PEAK | BL | 0,00 | 0,05 | 6978,00 | 0,00 | 1,0000 | 0,00 |
| ITBS - TACS | BL | 0,00 | 0,05 | 6978,00 | 0,00 | 1,0000 | 0,00 |
| X-TACS TROUGH - SHAM | 30 | 0,11 | 0,05 | 6979,10 | 2,19 | **0,0412** | 0,18 |
| X-TACS PEAK - SHAM | 30 | 0,31 | 0,05 | 6978,97 | 5,97 | **< 0.0001** | 0,51 |
| X-TACS PEAK - X-TACS TROUGH | 30 | 0,20 | 0,05 | 6978,53 | 3,91 | **0,0002** | 0,33 |
| TACS - SHAM | 30 | 0,26 | 0,05 | 6979,01 | 5,08 | **< 0.0001** | 0,42 |
| TACS - X-TACS TROUGH | 30 | 0,15 | 0,05 | 6978,10 | 2,97 | **0,0050** | 0,24 |
| TACS - X-TACS PEAK | 30 | -0,05 | 0,05 | 6978,58 | -0,99 | 0,3571 | 0,08 |
| ITBS - SHAM | 30 | 0,31 | 0,05 | 6978,88 | 6,19 | **< 0.0001** | 0,52 |
| ITBS - X-TACS TROUGH | 30 | 0,20 | 0,05 | 6978,33 | 4,12 | **0,0001** | 0,34 |
| ITBS - X-TACS PEAK | 30 | 0,01 | 0,05 | 6978,59 | 0,17 | 0,8676 | 0,01 |
| ITBS - TACS | 30 | 0,06 | 0,05 | 6978,33 | 1,17 | 0,3019 | 0,10 |
| X-TACS TROUGH - SHAM | 60 | 0,15 | 0,05 | 6978,32 | 2,95 | **0,0052** | 0,24 |
| X-TACS PEAK - SHAM | 60 | 0,33 | 0,05 | 6978,79 | 6,43 | **< 0.0001** | 0,54 |
| X-TACS PEAK - X-TACS TROUGH | 60 | 0,18 | 0,05 | 6978,53 | 3,53 | **0,0014** | 0,29 |
| TACS - SHAM | 60 | 0,08 | 0,05 | 6978,81 | 1,54 | 0,1534 | 0,12 |
| TACS - X-TACS TROUGH | 60 | -0,07 | 0,05 | 6978,73 | -1,39 | 0,1844 | 0.11 |
| TACS - X-TACS PEAK | 60 | -0,25 | 0,05 | 6979,26 | -4,86 | **< 0.0001** | 0.41 |
| ITBS - SHAM | 60 | 0,16 | 0,05 | 6978,33 | 3,19 | **0,0028** | 0,26 |
| ITBS - X-TACS TROUGH | 60 | 0,01 | 0,05 | 6978,30 | 0,23 | 0,8151 | 0,02 |
| ITBS - X-TACS PEAK | 60 | -0,17 | 0,05 | 6978,68 | -3,31 | **0,0023** | 0,28 |
| ITBS - TACS | 60 | 0,08 | 0,05 | 6978,80 | 1,62 | 0,1505 | 0,13 |
| X-TACS TROUGH - SHAM | 90 | 0,18 | 0,05 | 6978,84 | 3,50 | **0,0007** | 0,29 |
| X-TACS PEAK - SHAM | 90 | 0,41 | 0,05 | 6979,64 | 7,97 | **< 0.0001** | 0,68 |
| X-TACS PEAK - X-TACS TROUGH | 90 | 0,23 | 0,05 | 6978,94 | 4,51 | **< 0.0001** | 0,39 |
| TACS - SHAM | 90 | 0,44 | 0,05 | 6978,63 | 8,80 | **< 0.0001** | 0,73 |
| TACS - X-TACS TROUGH | 90 | 0,26 | 0,05 | 6978,74 | 5,26 | **< 0.0001** | 0,44 |
| TACS - X-TACS PEAK | 90 | 0,03 | 0,05 | 6979,55 | 0,63 | 0,5286 | 0,05 |
| ITBS - SHAM | 90 | 0,23 | 0,05 | 6978,47 | 4,68 | **0,000** | 0,39 |
| ITBS - X-TACS TROUGH | 90 | 0,06 | 0,05 | 6978,58 | 1,13 | 0,2892 | 0,09 |
| ITBS - X-TACS PEAK | 90 | -0,18 | 0,05 | 6979,39 | -3,46 | **0,0007** | 0,29 |
| ITBS - TACS | 90 | -0,21 | 0,05 | 6978,26 | -4,19 | **< 0.0001** | 0,34 |
| X-TACS TROUGH - SHAM | 120 | 0,22 | 0,05 | 6978,55 | 4,46 | **< 0.0001** | 0,37 |
| X-TACS PEAK - SHAM | 120 | 0,37 | 0,05 | 6978,90 | 7,27 | **< 0.0001** | 0,62 |
| X-TACS PEAK - X-TACS TROUGH | 120 | 0,15 | 0,05 | 6978,40 | 2,92 | **0,0070** | 0,24 |
| TACS - SHAM | 120 | 0,27 | 0,05 | 6978,70 | 5,32 | **< 0.0001** | 0,45 |
| TACS - X-TACS TROUGH | 120 | 0,05 | 0,05 | 6978,52 | 0,92 | 0,4502 | 0,07 |
| TACS - X-TACS PEAK | 120 | -0,10 | 0,05 | 6979,10 | -1,99 | 0,0668 | 0,16 |
| ITBS - SHAM | 120 | 0,24 | 0,05 | 6978,63 | 4,82 | **< 0.0001** | 0,40 |
| ITBS - X-TACS TROUGH | 120 | 0,02 | 0,05 | 6978,42 | 0,39 | 0,6979 | 0,03 |
| ITBS - X-TACS PEAK | 120 | -0,13 | 0,05 | 6978,98 | -2,52 | **0,0196** | 0,21 |
| ITBS - TACS | 120 | -0,03 | 0,05 | 6978,45 | -0,53 | 0,6659 | 0,04 |

**Table S5:** Pairwise comparisons between time windows 40Hz eyes-open

| **contrast** | **Condition** | **estimate** | **SE** | **df** | **t-value** | p-value | | d |
| --- | --- | --- | --- | --- | --- | --- | --- | --- |
| 30 - BL | SHAM | -0,03 | 0,05 | 6978,99 | -0,60 | | 0,6062 | 0,05 |
| 60 - BL | SHAM | 0,08 | 0,05 | 6978,17 | 1,65 | | 0,1661 | 0,13 |
| 60 - 30 | SHAM | 0,11 | 0,05 | 6978,93 | 2,20 | | 0,0928 | 0,18 |
| 90 - BL | SHAM | 0,06 | 0,05 | 6978,36 | 1,13 | | 0,3210 | 0,09 |
| 90 - 30 | SHAM | 0,09 | 0,05 | 6978,63 | 1,70 | | 0,1661 | 0,14 |
| 90 - 60 | SHAM | -0,02 | 0,05 | 6978,44 | -0,49 | | 0,6255 | 0,04 |
| 120 - BL | SHAM | 0,15 | 0,05 | 6978,40 | 3,08 | | **0,0103** | 0,25 |
| 120 - 30 | SHAM | 0,18 | 0,05 | 6978,63 | 3,59 | | **0,0033** | 0,31 |
| 120 - 60 | SHAM | 0,07 | 0,05 | 6978,40 | 1,45 | | 0,2116 | 0,12 |
| 120 - 90 | SHAM | 0,10 | 0,05 | 6978,24 | 1,91 | | 0,1399 | 0,16 |
| 30 - BL | X-TACS TROUGH | 0,08 | 0,05 | 6978,07 | 1,63 | | 0,1145 | 0,13 |
| 60 - BL | X-TACS TROUGH | 0,23 | 0,05 | 6978,13 | 4,63 | | **< 0.0001** | 0,38 |
| 60 - 30 | X-TACS TROUGH | 0,15 | 0,05 | 6978,18 | 3,00 | | **0,0039** | 0,25 |
| 90 - BL | X-TACS TROUGH | 0,23 | 0,05 | 6978,54 | 4,71 | | **< 0.0001** | 0,39 |
| 90 - 30 | X-TACS TROUGH | 0,15 | 0,05 | 6978,50 | 3,10 | | **0,0039** | 0,26 |
| 90 - 60 | X-TACS TROUGH | 0,01 | 0,05 | 6978,32 | 0,14 | | 0,8922 | 0,01 |
| 120 - BL | X-TACS TROUGH | 0,38 | 0,05 | 6978,12 | 7,72 | | **< 0.0001** | 0,62 |
| 120 - 30 | X-TACS TROUGH | 0,30 | 0,05 | 6978,17 | 6,06 | | **< 0.0001** | 0,49 |
| 120 - 60 | X-TACS TROUGH | 0,15 | 0,05 | 6978,07 | 3,04 | | **0,0039** | 0,25 |
| 120 - 90 | X-TACS TROUGH | 0,14 | 0,05 | 6978,22 | 2,86 | | **0,0052** | 0,24 |
| 30 - BL | X-TACS PEAK | 0,28 | 0,05 | 6978,47 | 5,52 | | **< 0.0001** | 0,46 |
| 60 - BL | X-TACS PEAK | 0,41 | 0,05 | 6978,64 | 8,11 | | **< 0.0001** | 0,51 |
| 60 - 30 | X-TACS PEAK | 0,13 | 0,05 | 6978,30 | 2,55 | | **0,0153** | 0,22 |
| 90 - BL | X-TACS PEAK | 0,47 | 0,05 | 6979,19 | 9,26 | | **< 0.0001** | 0,78 |
| 90 - 30 | X-TACS PEAK | 0,19 | 0,05 | 6978,76 | 3,70 | | **0,0004** | 0,32 |
| 90 - 60 | X-TACS PEAK | 0,06 | 0,05 | 6978,27 | 1,15 | | 0,2626 | 0,10 |
| 120 - BL | X-TACS PEAK | 0,52 | 0,05 | 6978,51 | 10,56 | | **< 0.0001** | 0,87 |
| 120 - 30 | X-TACS PEAK | 0,25 | 0,05 | 6978,43 | 4,88 | | **< 0.0001** | 0,41 |
| 120 - 60 | X-TACS PEAK | 0,12 | 0,05 | 6978,17 | 2,29 | | **0,0272** | 0,20 |
| 120 - 90 | X-TACS PEAK | 0,06 | 0,05 | 6978,36 | 1,12 | | 0,2626 | 0,10 |
| 30 - BL | TACS | 0,23 | 0,05 | 6978,09 | 4,61 | | **< 0.0001** | 0,37 |
| 60 - BL | TACS | 0,16 | 0,05 | 6978,69 | 3,19 | | **0,0018** | 0,27 |
| 60 - 30 | TACS | -0,07 | 0,05 | 6978,71 | -1,35 | | 0,1763 | 0,11 |
| 90 - BL | TACS | 0,50 | 0,05 | 6978,29 | 10,15 | | **< 0.0001** | 0,83 |
| 90 - 30 | TACS | 0,27 | 0,05 | 6978,37 | 5,51 | | **< 0.0001** | 0,45 |
| 90 - 60 | TACS | 0,34 | 0,05 | 6978,17 | 6,80 | | **< 0.0001** | 0,57 |
| 120 - BL | TACS | 0,42 | 0,05 | 6978,46 | 8,56 | | **< 0.0001** | 0,70 |
| 120 - 30 | TACS | 0,20 | 0,05 | 6978,53 | 3,97 | | **< 0,0001** | 0,33 |
| 120 - 60 | TACS | 0,27 | 0,05 | 6978,12 | 5,27 | | **< 0.0001** | 0,44 |
| 120 - 90 | TACS | -0,08 | 0,05 | 6978,10 | -1,50 | | 0,1481 | 0,13 |
| 30 - BL | ITBS | 0,28 | 0,05 | 6978,30 | 5,75 | | **< 0.0001** | 0,47 |
| 60 - BL | ITBS | 0,24 | 0,05 | 6978,13 | 4,88 | | **< 0.0001** | 0,39 |
| 60 - 30 | ITBS | -0,04 | 0,05 | 6978,31 | -0,90 | | 0,4096 | 0,07 |
| 90 - BL | ITBS | 0,29 | 0,05 | 6978,07 | 5,93 | | **< 0.0001** | 0,48 |
| 90 - 30 | ITBS | 0,01 | 0,05 | 6978,31 | 0,14 | | 0,8897 | 0,01 |
| 90 - 60 | ITBS | 0,05 | 0,05 | 6978,14 | 1,04 | | 0,3702 | 0,09 |
| 120 - BL | ITBS | 0,40 | 0,05 | 6978,34 | 8,06 | | **< 0.0001** | 0.66 |
| 120 - 30 | ITBS | 0,11 | 0,05 | 6978,16 | 2,26 | | **0,0395** | 0,19 |
| 120 - 60 | ITBS | 0,16 | 0,05 | 6978,39 | 3,18 | | **0,0030** | 0,26 |
| 120 - 90 | ITBS | 0,11 | 0,05 | 6978,36 | 2,14 | | **0,0465** | 0,18 |

**Table S6**: Pairwise comparisons between conditions 40Hz eyes-closed

| **contrast** | **Time** | **estimate** | **SE** | **df** | **t-value** | **p-value** |  | **d** |
| --- | --- | --- | --- | --- | --- | --- | --- | --- |
| X-TACS TROUGH - SHAM | BL | 0,00 | 0,04 | 7111,00 | 0,00 | 1,0000 |  | 0,00 |
| X-TACS PEAK - SHAM | BL | 0,00 | 0,04 | 7111,00 | 0,00 | 1,0000 |  | 0,00 |
| X-TACS Peak - X-TACS TROUGH | BL | 0,00 | 0,04 | 7111,00 | 0,00 | 1,0000 |  | 0,00 |
| TACS - SHAM | BL | 0,00 | 0,04 | 7111,00 | 0,00 | 1,0000 |  | 0,00 |
| TACS - X-TACS TROUGH | BL | 0,00 | 0,04 | 7111,00 | 0,00 | 1,0000 |  | 0,00 |
| TACS - X-TACS PEAK | BL | 0,00 | 0,04 | 7111,00 | 0,00 | 1,0000 |  | 0,00 |
| ITBS - SHAM | BL | 0,00 | 0,04 | 7111,00 | 0,00 | 1,0000 |  | 0,00 |
| ITBS - X-TACS TROUGH | BL | 0,00 | 0,04 | 7111,00 | 0,00 | 1,0000 |  | 0,00 |
| ITBS - X-TACS PEAK | BL | 0,00 | 0,04 | 7111,00 | 0,00 | 1,0000 |  | 0,00 |
| ITBS - TACS | BL | 0,00 | 0,04 | 7111,00 | 0,00 | 1,0000 |  | 0,00 |
| X-TACS TROUGH - SHAM | 30 | -0,16 | 0,04 | 7111,41 | -3,76 | **0,0017** |  | 0,31 |
| X-TACS PEAK - SHAM | 30 | -0,08 | 0,04 | 7111,09 | -1,83 | 0,1342 |  | 0,15 |
| X-TACS PEAK - X-TACS TROUGH | 30 | 0,08 | 0,04 | 7111,43 | 1,95 | 0,1268 |  | 0,16 |
| TACS - SHAM | 30 | -0,02 | 0,04 | 7111,08 | -0,53 | 0,5969 |  | 0,04 |
| TACS - X-TACS TROUGH | 30 | 0,14 | 0,04 | 7111,41 | 3,25 | **0,0059** |  | 0,27 |
| TACS - X-TACS PEAK | 30 | 0,06 | 0,04 | 7111,08 | 1,31 | 0,3185 |  | 0,11 |
| ITBS - SHAM | 30 | -0,05 | 0,04 | 7111,13 | -1,18 | 0,3378 |  | 0,10 |
| ITBS - X-TACS TROUGH | 30 | 0,11 | 0,04 | 7111,43 | 2,58 | **0,0326** |  | 0,21 |
| ITBS - X-TACS PEAK | 30 | 0,03 | 0,04 | 7111,04 | 0,64 | 0,5773 |  | 0,05 |
| ITBS - TACS | 30 | -0,03 | 0,04 | 7111,11 | -0,66 | 0,5773 |  | 0,05 |
| X-TACS TROUGH - SHAM | 60 | -0,10 | 0,04 | 7111,19 | -2,41 | **0,0321** |  | 0,20 |
| X-TACS PEAK - SHAM | 60 | -0,02 | 0,04 | 7111,09 | -0,54 | 0,7396 |  | 0,04 |
| X-TACS PEAK - X-TACS TROUGH | 60 | 0,08 | 0,04 | 7111,23 | 1,87 | 0,0886 |  | 0,15 |
| TACS - SHAM | 60 | 0,11 | 0,04 | 7111,19 | 2,57 | **0,0253** |  | 0,21 |
| TACS - X-TACS TROUGH | 60 | 0,21 | 0,04 | 7111,29 | 4,98 | **< 0.0001** |  | 0,41 |
| TACS - X-TACS PEAK | 60 | 0,13 | 0,04 | 7111,23 | 3,10 | **0,0096** |  | 0,26 |
| ITBS - SHAM | 60 | -0,01 | 0,04 | 7111,18 | -0,29 | 0,8045 |  | 0,02 |
| ITBS - X-TACS TROUGH | 60 | 0,09 | 0,04 | 7111,26 | 2,12 | 0,0574 |  | 0,17 |
| ITBS - X-TACS PEAK | 60 | 0,01 | 0,04 | 7111,26 | 0,25 | 0,8045 |  | 0,02 |
| ITBS - TACS | 60 | -0,12 | 0,04 | 7111,28 | -2,86 | **0,0143** |  | 0,24 |
| X-TACS TROUGH - SHAM | 90 | -0,18 | 0,04 | 7111,48 | -4,22 | **0,0002** |  | 0,35 |
| X-TACS PEAK - SHAM | 90 | -0,14 | 0,04 | 7111,53 | -3,18 | **0,0058** |  | 0,27 |
| X-TACS PEAK - X-TACS TROUGH | 90 | 0,04 | 0,04 | 7111,58 | 1,02 | 0,3399 |  | 0,09 |
| TACS - SHAM | 90 | -0,05 | 0,04 | 7111,26 | -1,21 | 0,3112 |  | 0,10 |
| TACS - X-TACS TROUGH | 90 | 0,13 | 0,04 | 7111,13 | 3,08 | **0,0058** |  | 0.25 |
| TACS - X-TACS PEAK | 90 | 0,09 | 0,04 | 7111,45 | 2,02 | 0,0862 |  | 0.17 |
| ITBS - SHAM | 90 | -0,13 | 0,04 | 7111,29 | -3,05 | **0,0058** |  | 0.26 |
| ITBS - X-TACS TROUGH | 90 | 0,05 | 0,04 | 7111,30 | 1,15 | 0,3112 |  | 0,10 |
| ITBS - X-TACS PEAK | 90 | 0,01 | 0,04 | 7111,40 | 0,13 | 0,8971 |  | 0,01 |
| ITBS - TACS | 90 | -0,08 | 0,04 | 7111,22 | -1,89 | 0,0981 |  | 0,16 |
| X-TACS TROUGH - SHAM | 120 | -0,20 | 0,04 | 7111,35 | -4,60 | **< 0.0001** |  | 0,38 |
| X-TACS PEAK - SHAM | 120 | -0,14 | 0,04 | 7111,61 | -3,16 | **0,0040** |  | 0,26 |
| X-TACS PEAK - X-TACS TROUGH | 120 | 0,06 | 0,04 | 7111,69 | 1,41 | 0,2282 |  | 0,12 |
| TACS - SHAM | 120 | 0,00 | 0,04 | 7111,11 | -0,10 | 0,9164 |  | 0,01 |
| TACS - X-TACS TROUGH | 120 | 0,19 | 0,04 | 7111,24 | 4,53 | **< 0.0001** |  | 0,37 |
| TACS - X-TACS PEAK | 120 | 0,13 | 0,04 | 7111,68 | 3,08 | **0,0042** |  | 0,26 |
| ITBS - SHAM | 120 | -0,03 | 0,04 | 7111,25 | -0,67 | 0,6285 |  | 0,06 |
| ITBS - X-TACS TROUGH | 120 | 0,17 | 0,04 | 7111,32 | 3,93 | **0,0003** |  | 0,32 |
| ITBS - X-TACS PEAK | 120 | 0,11 | 0,04 | 7111,67 | 2,49 | **0,0212** |  | 0,20 |
| ITBS - TACS | 120 | -0,02 | 0,04 | 7111,12 | -0,57 | 0,6310 |  | 0,05 |

**Table S7:** Pairwise comparisons between time windows 40Hz eyes-closed

| **contrast** | **Condition** | **estimate** | **SE** | **df** | **t-value** | **p-value** | d |
| --- | --- | --- | --- | --- | --- | --- | --- |
| 30 - BL | SHAM | 0,22 | 0,04 | 7111,07 | 5,25 | **< 0.0001** | 0,43 |
| 60 - BL | SHAM | 0,25 | 0,04 | 7111,07 | 5,91 | **< 0.0001** | 0,48 |
| 60 - 30 | SHAM | 0,03 | 0,04 | 7111,05 | 0,66 | 0,5643 | 0,05 |
| 90 - BL | SHAM | 0,43 | 0,04 | 7111,35 | 9,92 | **< 0.0001** | 0,81 |
| 90 - 30 | SHAM | 0,20 | 0,04 | 7111,23 | 4,72 | **< 0.0001** | 0,39 |
| 90 - 60 | SHAM | 0,18 | 0,04 | 7111,13 | 4,05 | **0,0001** | 0,34 |
| 120 - BL | SHAM | 0,45 | 0,04 | 7111,11 | 10,58 | **< 0.0001** | 0,87 |
| 120 - 30 | SHAM | 0,23 | 0,04 | 7111,09 | 5,32 | **< 0.0001** | 0,44 |
| 120 - 60 | SHAM | 0,20 | 0,04 | 7111,04 | 4,65 | **< 0.0001** | 0,38 |
| 120 - 90 | SHAM | 0,02 | 0,04 | 7111,15 | 0,56 | 0,5775 | 0,05 |
| 30 - BL | X-TACS TROUGH | 0,06 | 0,04 | 7111,36 | 1,41 | 0,1754 | 0,12 |
| 60 - BL | X-TACS TROUGH | 0,15 | 0,04 | 7111,12 | 3,48 | **0,0010** | 0,28 |
| 60 - 30 | X-TACS TROUGH | 0,09 | 0,04 | 7111,16 | 2,02 | 0,0546 | 0,17 |
| 90 - BL | X-TACS TROUGH | 0,24 | 0,04 | 7111,11 | 5,72 | **< 0.0001** | 0,47 |
| 90 - 30 | X-TACS TROUGH | 0,18 | 0,04 | 7111,35 | 4,22 | **0,0001** | 0,35 |
| 90 - 60 | X-TACS TROUGH | 0,10 | 0,04 | 7111,21 | 2,23 | **0,0372** | 0,18 |
| 120 - BL | X-TACS TROUGH | 0,25 | 0,04 | 7111,17 | 5,90 | **< 0.0001** | 0,48 |
| 120 - 30 | X-TACS TROUGH | 0,19 | 0,04 | 7111,48 | 4,40 | **< 0.0001** | 0,37 |
| 120 - 60 | X-TACS TROUGH | 0,10 | 0,04 | 7111,27 | 2,42 | **0,0259** | 0,20 |
| 120 - 90 | X-TACS TROUGH | 0,01 | 0,04 | 7111,11 | 0,21 | 0,8373 | 0,01 |
| 30 - BL | X-TACS PEAK | 0,14 | 0,04 | 7111,04 | 3,42 | **0,0012** | 0,28 |
| 60 - BL | X-TACS PEAK | 0,23 | 0,04 | 7111,09 | 5,35 | **< 0.0001** | 0,44 |
| 60 - 30 | X-TACS PEAK | 0,08 | 0,04 | 7111,09 | 1,95 | 0,0646 | 0,16 |
| 90 - BL | X-TACS PEAK | 0,29 | 0,04 | 7111,47 | 6,71 | **< 0.0001** | 0,55 |
| 90 - 30 | X-TACS PEAK | 0,14 | 0,04 | 7111,50 | 3,32 | **0,0015** | 0,27 |
| 90 - 60 | X-TACS PEAK | 0,06 | 0,04 | 7111,31 | 1,38 | 0,1872 | 0,11 |
| 120 - BL | X-TACS PEAK | 0,31 | 0,04 | 7111,60 | 7,28 | **< 0.0001** | 0,60 |
| 120 - 30 | X-TACS PEAK | 0,17 | 0,04 | 7111,45 | 3,91 | **0,0002** | 0,32 |
| 120 - 60 | X-TACS PEAK | 0,09 | 0,04 | 7111,28 | 1,97 | 0,0646 | 0,16 |
| 120 - 90 | X-TACS PEAK | 0,03 | 0,04 | 7111,63 | 0,59 | 0,5535 | 0,05 |
| 30 - BL | TACS | 0,20 | 0,04 | 7111,03 | 4,74 | **< 0.0001** | 0,39 |
| 60 - BL | TACS | 0,36 | 0,04 | 7111,14 | 8,51 | **< 0.0001** | 0,69 |
| 60 - 30 | TACS | 0,16 | 0,04 | 7111,14 | 3,78 | **0,0002** | 0,31 |
| 90 - BL | TACS | 0,37 | 0,04 | 7111,02 | 8,87 | **< 0.0001** | 0,71 |
| 90 - 30 | TACS | 0,17 | 0,04 | 7111,04 | 4,12 | **0,0001** | 0,33 |
| 90 - 60 | TACS | 0,01 | 0,04 | 7111,16 | 0,31 | 0,7537 | 0,02 |
| 120 - BL | TACS | 0,44 | 0,04 | 7111,04 | 10,57 | **< 0.0001** | 0,86 |
| 120 - 30 | TACS | 0,25 | 0,04 | 7111,06 | 5,81 | **< 0.0001** | 0,47 |
| 120 - 60 | TACS | 0,09 | 0,04 | 7111,17 | 2,00 | 0,0565 | 0,16 |
| 120 - 90 | TACS | 0,07 | 0,04 | 7111,03 | 1,70 | 0,0991 | 0,14 |
| 30 - BL | ITBS | 0,17 | 0,04 | 7111,07 | 4,06 | **0,0001** | 0,33 |
| 60 - BL | ITBS | 0,24 | 0,04 | 7111,13 | 5,60 | **< 0.0001** | 0,45 |
| 60 - 30 | ITBS | 0,07 | 0,04 | 7111,14 | 1,55 | 0,1345 | 0,13 |
| 90 - BL | ITBS | 0,29 | 0,04 | 7111,27 | 6,83 | **< 0.0001** | 0,56 |
| 90 - 30 | ITBS | 0,12 | 0,04 | 7111,24 | 2,80 | **0,0063** | 0,23 |
| 90 - 60 | ITBS | 0,05 | 0,04 | 7111,18 | 1,26 | 0,2075 | 0,10 |
| 120 - BL | ITBS | 0,42 | 0,04 | 7111,13 | 9,90 | **< 0.0001** | 0,81 |
| 120 - 30 | ITBS | 0,25 | 0,04 | 7111,08 | 5,82 | **< 0.0001** | 0,48 |
| 120 - 60 | ITBS | 0,18 | 0,04 | 7111,03 | 4,26 | **< 0.0001** | 0,35 |
| 120 - 90 | ITBS | 0,13 | 0,04 | 7111,18 | 2,97 | **0,0043** | 0,25 |
